# Spatially Distinct Myosin II Architectures Regulate Protrusion Dynamics and Directional Persistence during Immune Cell Migration

**DOI:** 10.64898/2026.03.13.711384

**Authors:** Nicolas Melis, Desu Chen, Emily Chen, Thomas Madsen, Yeap Ng, Bhagawat Subramanian, Weiye Wang, Carole Parent, Wolfgang Losert, Roberto Weigert

## Abstract

Directional persistence is essential for efficient immune cell migration in tissues, yet how cytoskeletal systems stabilize migration in complex three-dimensional environments remains unclear. Using intravital subcellular microscopy and quantitative analysis of membrane dynamics, we identify two spatially distinct architectures of non-muscle myosin II (NMII) that coordinate protrusion dynamics during neutrophil migration. In vivo and in collagen matrices, NMII assembles at the leading edge into lattice-like structures that are structurally and functionally distinct from rear contractile actomyosin bundles. Protrusion-resolved analyses reveal that directional persistence correlates strongly with protrusion lifetime and sustained NMII engagement, with rear NMII load showing the strongest association with protrusion persistence. Strikingly, directional migration is not determined by the abundance of favorable protrusions but by their temporal organization during migration. Pharmacological perturbations that redistribute NMII activity disrupt this temporal organization and alter migration trajectories. Together, these findings reveal that spatially distinct NMII architectures coordinate protrusion dynamics across time to stabilize directional migration in complex environments.

**Highlights:**

- Migrating neutrophils assemble spatially distinct myosin II architectures at the leading edge and rear
- Protrusion dynamics and directional persistence are linked to sustained myosin II engagement
- Directional migration emerges from the temporal organization of protrusion states rather than their abundance

## Introduction

Efficient migration of immune cells through tissues is essential for host defense, tissue repair, and immune surveillance^1–3^. To reach sites of infection, injury, or pathology, immune cells must navigate complex three-dimensional environments composed of heterogeneous extracellular matrices, densely packed cells, and dynamically evolving chemical and mechanical cues^4–6^. In these contexts, migration efficiency depends not only on speed but critically on the ability to maintain directional persistence over time^7^. Excessive turning or loss of polarity can delay arrival at target sites, reduce functional responses, and compromise coordinated immune behavior. Despite its importance, the mechanisms that stabilize directional persistence during immune cell migration in vivo remain incompletely understood.

Directional persistence is increasingly recognized as a regulated property of cell migration rather than a passive outcome of protrusion and contractility^7–9^. Migrating immune cells must dynamically balance exploratory behavior with sustained forward movement, adjusting their persistence in response to tissue architecture, chemotactic signals, and mechanical constraints^10,11^. While high persistence can accelerate targeting of localized insults, reduced persistence may facilitate tissue scanning or obstacle avoidance. Understanding how cytoskeletal systems regulate this balance is therefore central to immune cell biology.

Neutrophils provide a powerful model for studying immune cell migration in vivo. As rapid responders to infection and sterile injury, neutrophils extravasate from the vasculature and migrate through interstitial tissues to reach sites of damage^1,2,12^. Their migration is characterized by fast amoeboid movement, low adhesion to the extracellular matrix, and continuous remodeling of the plasma membrane through cycles of protrusion and retraction^13,14^. Maintaining directional persistence while navigating highly confined and mechanically heterogeneous environments is therefore a key requirement for effective neutrophil function.

Much of our current understanding of neutrophil migration derives from simplified two-dimensional systems, including under-agarose assays and microfabricated channels^15,16^. In these settings, migration is thought to arise from the coordinated activity of actin polymerization at the leading edge and non-muscle myosin II (NMII)– driven contractility at the rear of the cell^17,18^. Actin-based protrusions initiate forward movement, whereas rear-localized NMII generates contractile forces that retract the trailing edge and support forward translocation^18–20^. Although these models have provided important insights into cytoskeletal regulation of migration, they do not fully capture the structural and mechanical complexity of interstitial tissues.

Recent studies have revealed substantial differences between immune cell migration in two-dimensional environments and in three-dimensional tissues, including altered force transmission, cytoskeletal organization, and signaling dynamics^4,21,22^. In particular, how cytoskeletal systems are spatially organized and dynamically regulated in vivo remains poorly understood. Whether mechanisms identified in simplified systems fully explain directional persistence during immune migration in tissues therefore remains an open question.

NMII has traditionally been viewed as a rear-localized motor whose primary role during migration is to generate contractile forces that retract the trailing edge^11,15,23^. In neutrophils, NMIIA is the sole expressed NMII isoform and has been implicated in polarity maintenance, uropod retraction, and coordination of front–rear dynamics^18,24,25^. However, increasing evidence from diverse cellular contexts indicates that myosin II can adopt distinct supramolecular organizations and mechanical functions depending on its spatial localization and higher-order assembly^26–28^.

Whether NMII contributes directly to leading-edge dynamics during immune cell migration in vivo has remained largely unexplored. In particular, it is unclear whether myosin II participates in regulating protrusion stability, selection, or persistence in three-dimensional environments, or whether its functions are restricted to the trailing edge.

Here, using intravital subcellular microscopy in live mice combined with quantitative analysis of membrane and cytoskeletal dynamics, we uncover an unexpected spatial organization of NMII during neutrophil migration. We show that in vivo and in three-dimensional collagen matrices, NMII assembles not only at the rear of migrating cells but also at the leading edge, where it forms lattice-like structures that are structurally distinct from rear contractile actomyosin bundles. These observations reveal that NMII is organized into spatially specialized architectures during migration in complex environments.

Because the leading edge of migrating neutrophils is composed of dynamically generated protrusions, directional migration emerges from the coordinated behavior of transient protrusive events rather than from static front–rear polarity alone. By combining protrusion-resolved measurements with cell-scale migration analysis, we show that directional persistence is strongly linked to protrusion lifetime and sustained NMII engagement. Moreover, classification of protrusion states demonstrates that directional migration is not determined by the abundance of favorable protrusions but by their temporal organization during migration.

Together, these findings reveal that neutrophils deploy spatially distinct NMII architectures that coordinate protrusion dynamics across time to stabilize directional migration in complex environments. This work identifies a previously unrecognized mechanism through which spatial organization of the actomyosin cytoskeleton regulates directional persistence during immune cell migration in physiologically relevant tissues.

## Results

### Myosin II is recruited to the leading edge of migrating neutrophils in vivo and in 3D environments

To visualize the spatial distribution of NMII during neutrophil migration in vivo, we used an established laser-induced injury model in the mouse ear^3,19^. Neutrophils were purified from knock-in mice ubiquitously expressing GFP-NMIIA and adoptively transferred into the ears of recipient wild-type mice (Fig. 1A and S1A)^14,17,19^. Following laser injury, neutrophils rapidly accumulated and migrated through the collagen network toward the wound site with the characteristic amoeboid modality (Fig. 1B, asterisk; Movie S1). As previously described in two-dimensional migration systems, GFP-NMII was prominently enriched at the rear of migrating cells (Fig. 1B, red dashed outline)^10,12,15,28^. Unexpectedly, however, NMII was also consistently detected at the leading edge of migrating neutrophils (Fig. 1B, blue dashed outline; Movie S2).

**Figure 1.**
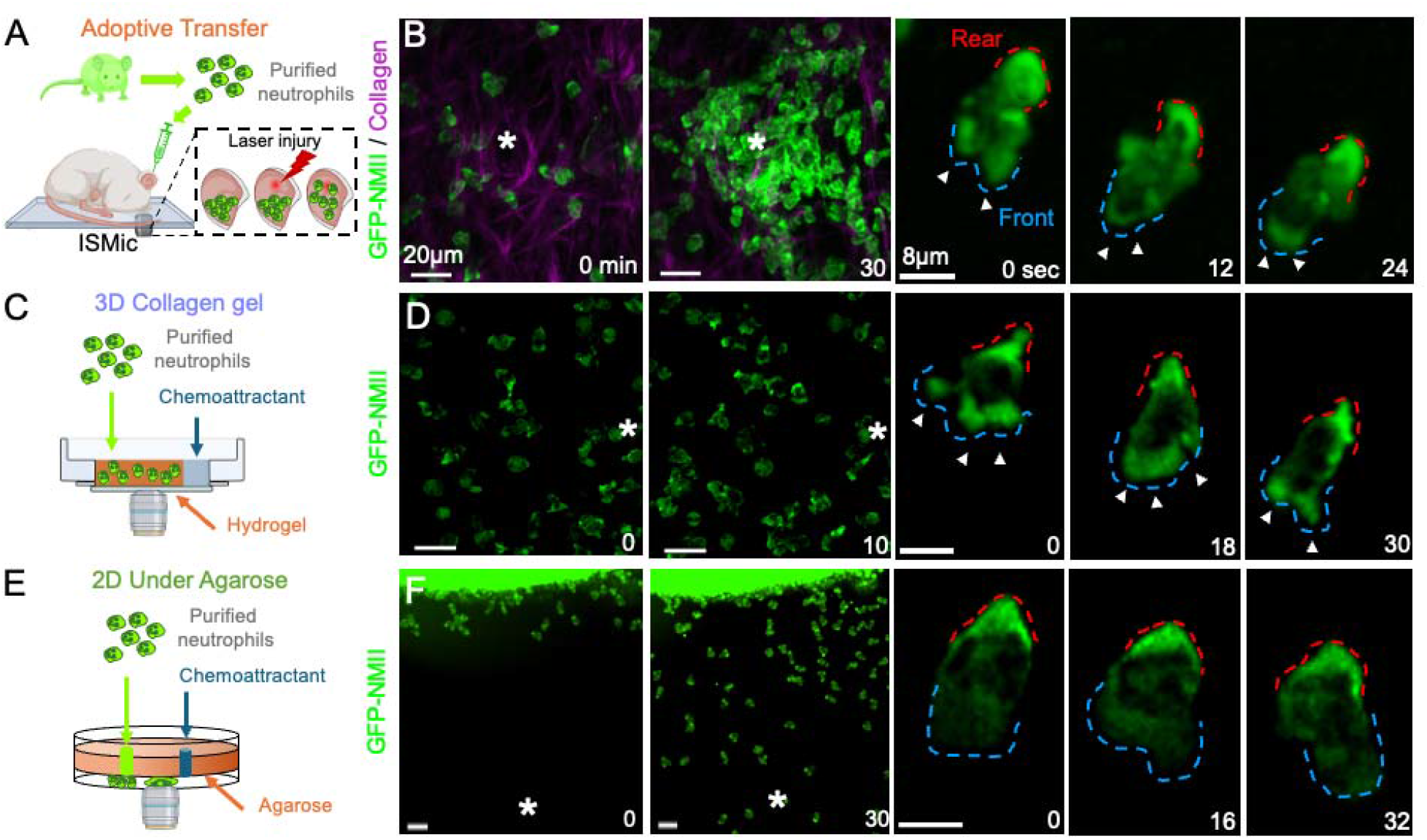
NMII localizes to the leading edge of migrating neutrophils in vivo and in three-dimensional environments. Primary neutrophils isolated from GFP-NMIIA knock-in mice were used to visualize NMII localization during migration in vivo and in vitro. (A) Schematic of the adoptive transfer approach used for intravital subcellular microscopy (ISMic), in which purified GFP-NMIIA neutrophils were injected into the ear dermis of recipient mice and imaged during migration toward a laser-induced injury site. (B) Two-photon imaging of neutrophils migrating in the mouse ear following laser injury (asterisk). Maximum-intensity projections show cells at 0 and 30 min after injury, with collagen fibers visualized by second harmonic generation (magenta) and GFP-NMIIA shown in green. Time-lapse insets show a representative migrating neutrophil highlighting NMII localization at the rear (red dashed outline) and at the leading edge (blue dashed outline). (C) Schematic of the three-dimensional collagen migration assay in which purified neutrophils were embedded in collagen I hydrogels and stimulated with a chemoattractant gradient. (D) Maximum-intensity projections of neutrophils migrating in collagen gels before and after chemoattractant addition, with time-lapse images showing NMII localization at both the leading edge and rear of migrating cells. (E) Schematic of the two-dimensional under-agarose migration assay used to generate a confined chemotactic gradient. (F) Maximum-intensity projections of neutrophils migrating under agarose toward the chemoattractant source, with time-lapse images showing NMII localization predominantly at the rear of the cell. See Movies S1, S7, and S8.

This distribution was not attributable to the GFP tag. Neutrophils expressing cytosolic GFP displayed no polarized enrichment (Fig. S1B), and immunolabeling of wild-type neutrophils confirmed the presence of endogenous NMII at the leading edge (Fig. S1C). Moreover, front localization of NMII was observed across multiple in vivo contexts in which neutrophil recruitment and migration were triggered by (i) laser-induced injury in the ear (Fig. S1D–F; Movies S3 and S4), (ii) injection of bacterial-derived particles into the footpad (Fig. S1G–H; Movie S5)^27^, or (iii) carcinogen-induced tumors in the mouse tongue (Fig. S1I–J; Movie S6) ^29^. These observations indicate that NMII recruitment to the leading edge represents a general feature of neutrophil migration in vivo.

To determine whether this spatial distribution depends on the physical environment, we compared NMII localization during neutrophil migration in three-dimensional collagen matrices and in two-dimensional assays^13,30^. When mouse or human neutrophils migrated within collagen I hydrogels (Fig. 1C), NMII was detected both at the leading edge and at the rear of the cell (Fig. 1D and Fig. S1J; Movie S7). In contrast, during migration in a 2D under-agarose assay, NMII localization was largely restricted to the rear of the cell (Fig. 1E–F; Movie S8), consistent with previous observations^31,32^. Importantly, the same rear-restricted distribution was observed in the presence of collagen I in two-dimensional conditions, indicating that matrix composition alone does not induce NMII recruitment to the leading edge^13^. Together, these results suggest that the three-dimensional environment plays a key role in promoting NMII localization at the leading edge of migrating neutrophils.

Quantification of NMII distribution along the cell boundary confirmed the qualitative observations described above. The average fraction of total cellular NMII localized at the leading edge, rear, and sides of migrating neutrophils was measured during migration in 2D, in 3D collagen matrices, and in vivo (Fig. 2A and Fig. S2A). In agreement with previous observations in two-dimensional systems, NMII was predominantly enriched at the rear of migrating cells in 2D conditions. In contrast, during migration in 3D collagen gels and in vivo, a substantial fraction of NMII was also detected at the leading edge (Fig. 2A). Time-lapse imaging revealed that NMII distribution at the leading edge was highly dynamic during migration. In individual cells, NMII levels at the front fluctuated substantially over time (Fig. 2B,C), whereas rear NMII levels were comparatively more stable. In some cells, NMII was transiently reduced or nearly absent at the leading edge during certain time intervals, indicating considerable heterogeneity in front NMII recruitment during migration.

**Figure 2.**
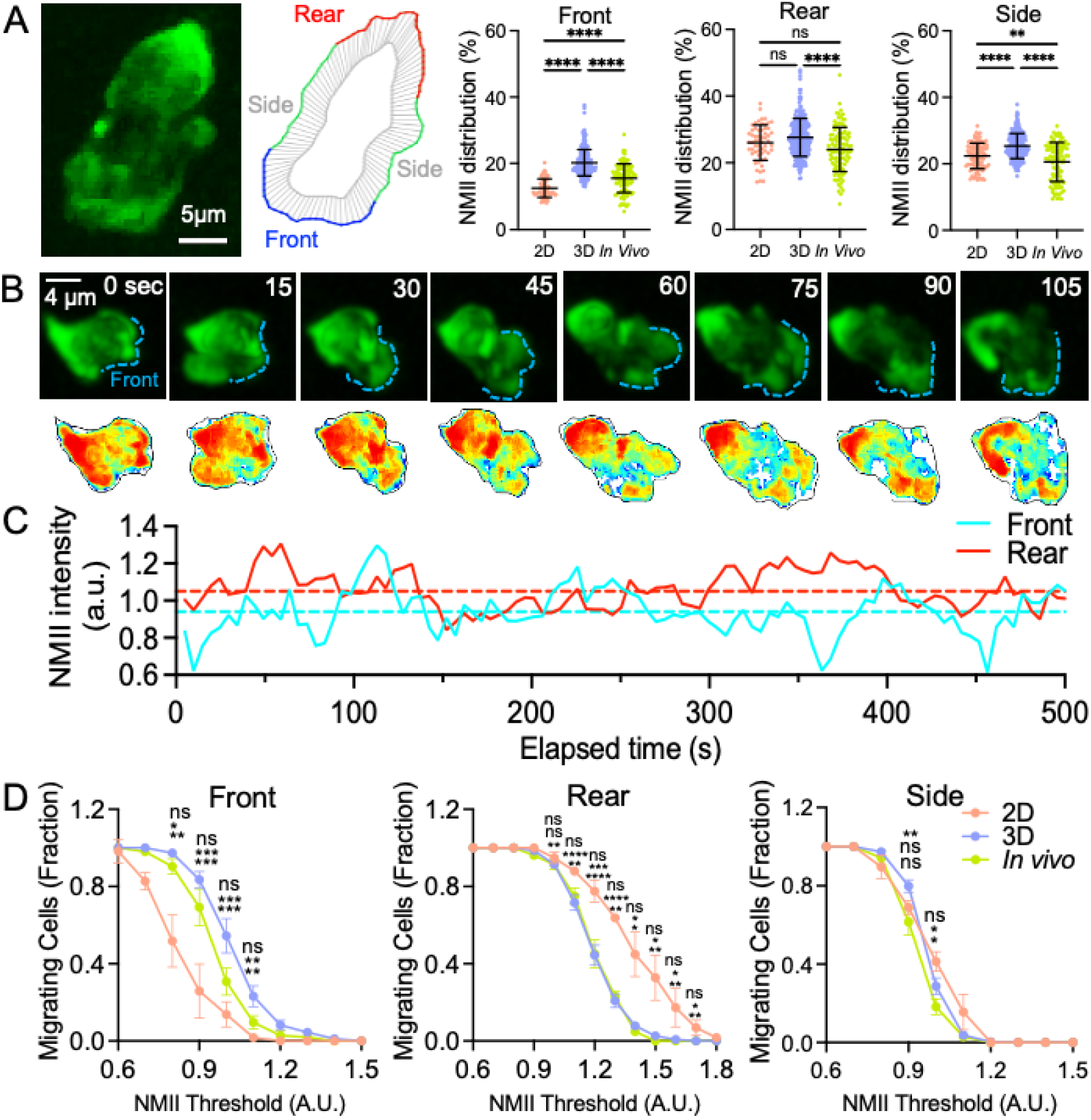
Dynamic distribution of NMII during neutrophil migration. (A) Quantification of GFP-NMII localization along the cell cortex during neutrophil migration in 2D, 3D collagen gels, and in vivo. The cell boundary was segmented into front, rear, and side regions as described in Methods, and the fraction of total cellular NMII localized to each region was calculated. Each dot represents an individual cell, and bars indicate mean ± SEM (2D: 58 cells, 3 experiments; 3D: 268 cells, 11 experiments; in vivo: 104 cells, 5 animals). Statistical significance was determined using one-way ANOVA with Tukey’s multiple comparisons test (**** p < 0.0001; ns, not significant). (B) Time-lapse imaging of GFP-NMII during neutrophil migration in vivo. Upper panels show maximum-intensity projections of a representative cell at selected time points, with the leading edge indicated by a blue dashed outline. Lower panels show corresponding heat maps representing the spatial distribution of NMII along the cell cortex. (C) Temporal fluctuations of normalized NMII intensity at the front (cyan) and rear (red) of the representative migrating cell shown in (B). Dashed lines indicate average NMII levels at the front and rear over the entire migration period. (D) Persistence analysis of NMII localization. The fraction of migrating cells maintaining NMII levels above increasing threshold values for more than 50% of the migration time was calculated separately for front, rear, and side regions in 2D, 3D, and in vivo conditions (see Methods). Data are shown as mean ± SEM of independent experiments.

To quantitatively assess this variability, we measured the fraction of cells in which NMII remained persistently enriched at a given region of the cell cortex. NMII localization was considered persistent when its level remained above a defined threshold for more than 50% of the migration time (see Methods). Across a range of NMII thresholds, a significantly larger fraction of neutrophils migrating in vivo or within collagen hydrogels exhibited persistent NMII enrichment at the leading edge compared with cells migrating in 2D (Fig. 2D). In contrast, persistent NMII enrichment at the rear was more prominent in cells migrating in 2D conditions. No major differences were observed in NMII localization at the sides of the cell among the different migration environments (Fig. 2D). Together, these results indicate that NMII recruitment to the leading edge is a dynamic feature of neutrophil migration in three-dimensional environments and that a substantial fraction of migrating cells maintain sustained NMII enrichment at the front during migration.

Similar results were obtained under more stringent conditions in which persistent NMII localization was defined as present for more than 67% of the migration time (Fig. S2B). At the spatial resolution of our imaging systems, we could not determine what fraction of NMII was cytoplasmic versus membrane-associated. Nonetheless, immunolabeling confirmed that NMII was in an active conformation, as its regulatory light chain was phosphorylated on Ser19, indicating competence for filament assembly and interaction with F-actin (Fig. S2C)^25,33^.

Because NMII displayed similar localization and dynamic behavior in neutrophils migrating in three-dimensional collagen matrices and in vivo, we next asked whether collagen hydrogels recapitulate additional aspects of neutrophil migration observed in living tissues. In both systems, neutrophils exhibited comparable morphology (Fig. S3A) and migrated with similar average speed (Fig. 3A) and directional persistence (Fig. 3B). The dynamic behaviors of the three plasma membrane domains defined in migrating neutrophils—front, rear, and side—were also similar between 3D hydrogels and in vivo conditions (see Methods and Fig. S4). In particular, the rates of leading-edge expansion and retraction were comparable in the two environments (Fig. 3C and Fig. S3B). To independently validate boundary measurements derived from GFP–NMIIA segmentation, neutrophils were labeled with the membrane dye MemGlow. MemGlow-based measurements of expansion and retraction rates closely matched those obtained using GFP–NMIIA segmentation, confirming that the latter accurately reflects plasma membrane dynamics (Fig. S3B). In both cases, front protrusion was temporally coupled to rear retraction, as indicated by the similar distributions of time offsets between maximal front expansion and the onset of rear contraction (Fig. 3F). Recruitment of NMII to the rear followed a similar temporal relationship with front protrusion dynamics (Fig. 3G). In addition, NMII localization was positively correlated with regions of positive membrane curvature at the cell front (Fig. 3D) and with area changes associated with rear retraction (Fig. 3E).

**Figure 3.**
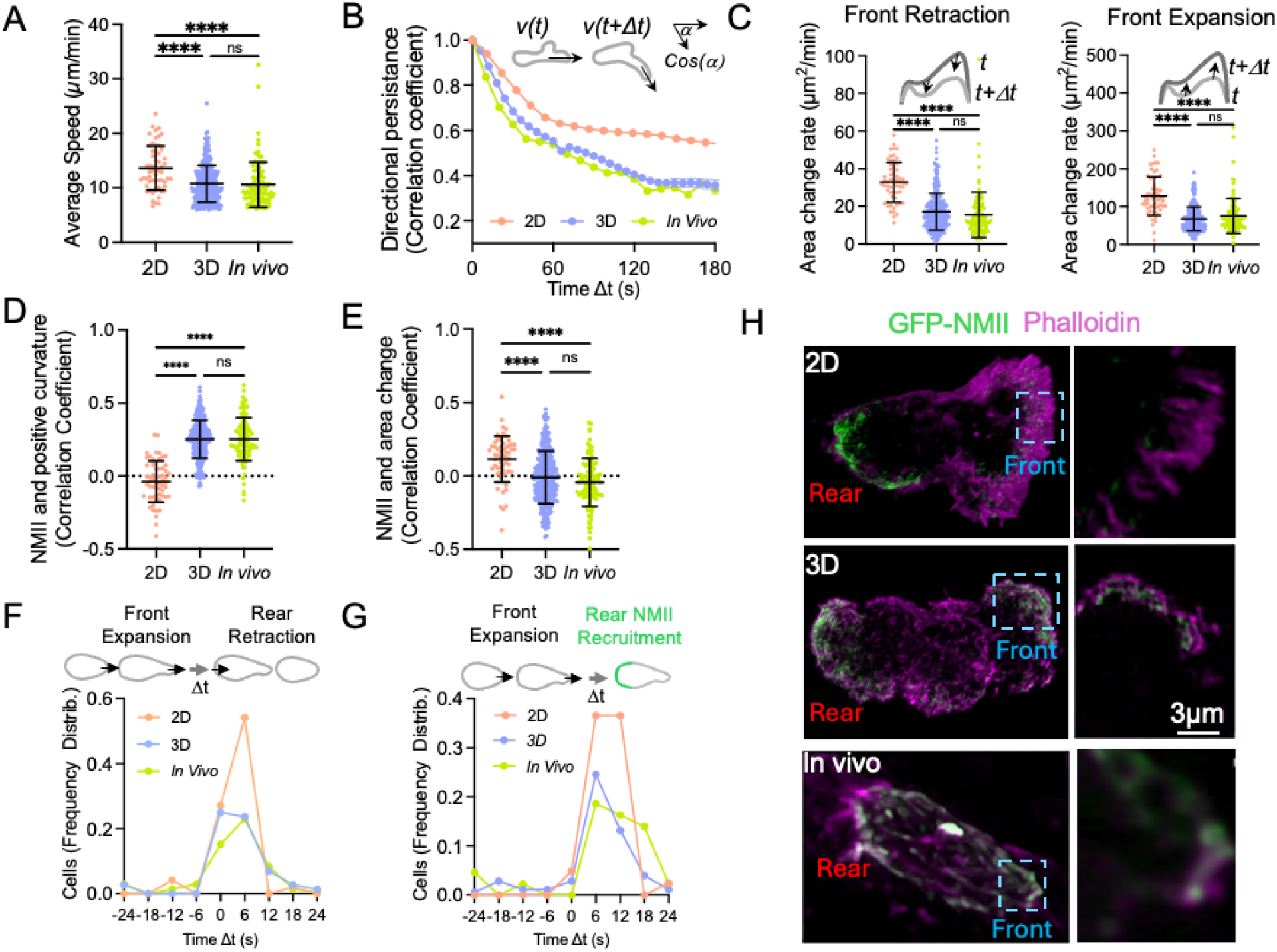
Collagen hydrogels recapitulate key features of neutrophil migration observed in vivo. Quantitative comparison of neutrophil migration in 2D under-agarose assays, 3D collagen gels, and in vivo conditions. (A) Average migration speed of neutrophils in each environment. (B) Directional persistence calculated as the autocorrelation of migration direction over increasing time offsets (Δt). (C) Rates of leading-edge retraction and expansion measured from boundary dynamics of migrating cells. (D) Correlation between local NMII levels and positive membrane curvature along the cell cortex. (E) Correlation between NMII localization and area changes associated with rear retraction. (F) Frequency distribution of time offsets between maximal front expansion and the onset of rear retraction. (G) Frequency distribution of time offsets between maximal front expansion and recruitment of NMII at the rear. (H) Representative images of neutrophils migrating in 2D, 3D collagen gels, or in vivo following fixation and labeling of GFP-NMII (green) and F-actin with phalloidin (magenta). Data in panels A, C, F, and G are presented as mean ± S.D. Statistical significance was determined using one-way ANOVA with Tukey’s multiple comparisons test (**p < 0.01, ****p < 0.0001).

We further examined the behavior of submicron protrusive structures that continuously form and retract at the leading edge (pseudopods^34,35^). Their frequency (Fig. S3D) and rates of assembly and disassembly (Fig. S3F–G) were comparable in 3D matrices and in vivo, although modest differences in protrusion lifetime were detected (Fig. S3E). Finally, cortical actin displayed a similar spatial distribution in 3D matrices and in vivo, whereas in two-dimensional migration actin was strongly enriched at the leading edge (Fig. 3H).

Together, these results demonstrate that collagen I hydrogels provide a tractable experimental system that faithfully reproduces key morphological, dynamic, and mechanical features of neutrophil migration observed in vivo, while enabling quantitative analysis of cytoskeletal dynamics that is difficult to achieve during intravital imaging. Having established that neutrophil migration in 3D hydrogels closely reproduces the dynamics observed in vivo, we next used this system to quantitatively analyze how protrusion behavior and NMII organization contribute to directional migration.

### Protrusion persistence links NMII polarity to directional migration

The presence of NMII at the leading edge raised the question of how this front-associated pool contributes to directional migration in three-dimensional environments. The leading edge of migrating neutrophils generates dynamic membrane protrusions, or pseudopods^34,36^, which extend, stabilize, and retract in a coordinated sequence. Directional migration therefore emerges not from static front–rear polarity alone but from the integrated behavior of transient protrusive events.

To characterize these dynamics, we resolved protrusions into three sequential phases: expansion (E), stabilization (S), and retraction (R) (Fig. 4A). Directional parameters were calculated across the lifetime of each protrusion, linking protrusion behavior directly to cell-scale migration.

**Figure 4.**
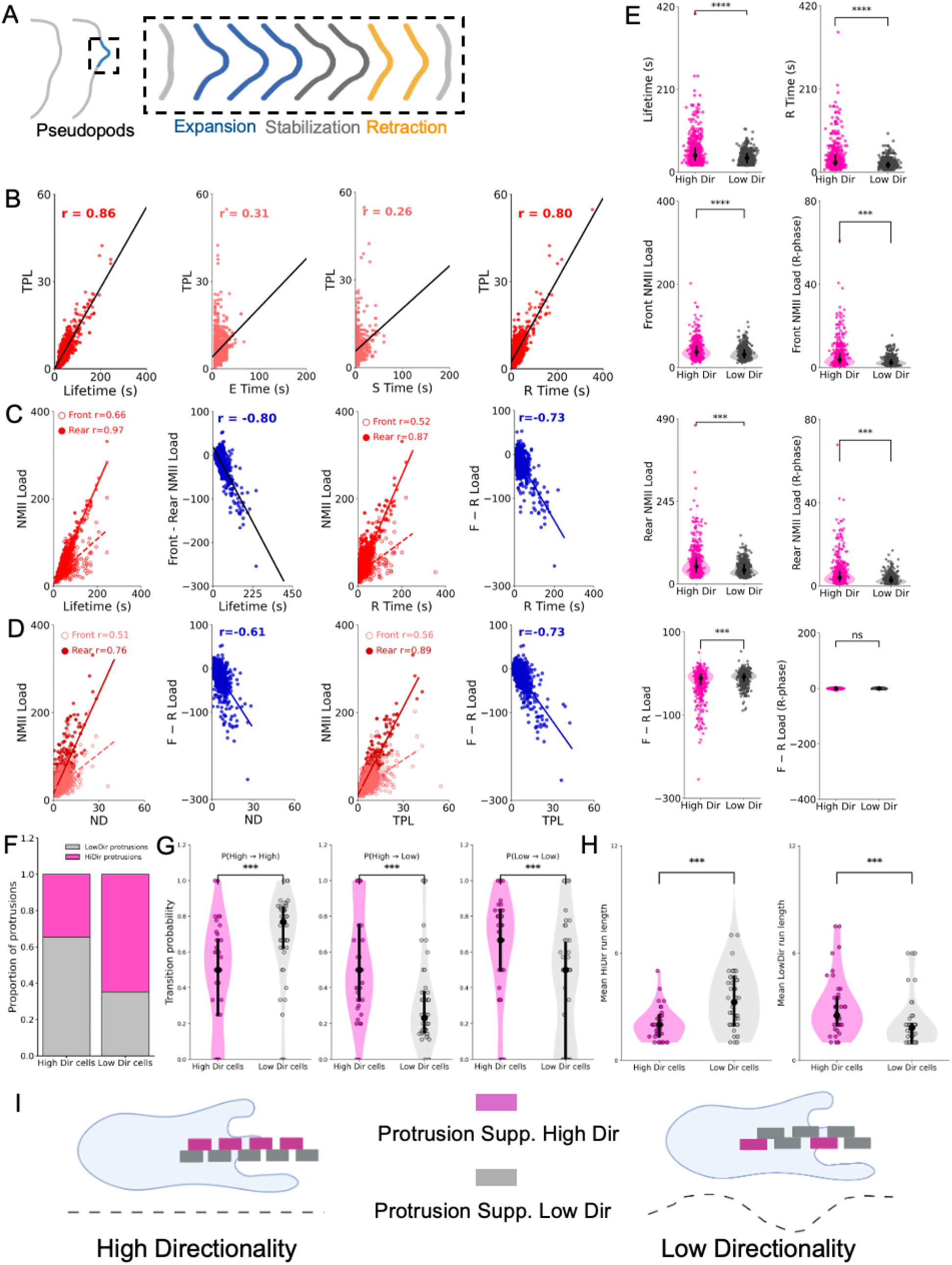
Protrusion persistence and NMII polarity determine directional migration. (A) Schematic representation of protrusion dynamics resolved into three phases: expansion (E), stabilization (S), and retraction (R), whose combined duration defines protrusion lifetime. (B) Correlations between protrusion structural parameters and directional migration measured as total path length (TPL), showing strong associations with protrusion lifetime and retraction duration. (C) Relationships between NMII engagement and protrusion persistence. Rear NMII load shows the strongest association with protrusion lifetime and retraction duration, whereas front NMII load exhibits weaker correlations. Front–rear polarity metrics display inverse relationships with protrusion lifetime. (D) Correlations between NMII metrics and net cell displacement (ND), indicating that rear NMII engagement is positively associated with directional migration, whereas front–rear dominance metrics show inverse associations (see also Fig. S4C). (E) Stratification of protrusions into high- and low-directionality groups using a composite directionality index derived from principal component analysis (see Methods; Fig. S5A–D). High-directionality protrusions exhibit longer lifetimes, extended retraction phases, and increased rear NMII engagement compared with low-directionality protrusions. Data are derived from 1,153 protrusions across 106 cells from independent experiments. Statistical comparisons were performed using nonparametric tests as described in Methods. (E) Stratification of protrusions into high- and low-directionality groups using a composite directionality index derived from principal component analysis (see Methods; Fig. S5A–D). High-directionality protrusions exhibit longer lifetimes, extended retraction phases, and increased rear NMII engagement compared with low-directionality protrusions. (F) Relative proportion of directionality-supporting and non– directionality-supporting protrusions in cells classified as highly directional or weakly directional. (G) Transition probabilities between protrusion states, showing that highly directional cells display increased transitions between states whereas low-directionality cells show higher probabilities of repeated transitions within the same state. (H) Distribution of protrusion run lengths in highly directional and low-directionality cells, indicating that highly directional cells exhibit shorter runs of similar protrusion states, whereas low-directionality cells display longer consecutive runs. (I) Conceptual model illustrating how directional migration emerges from the temporal organization of protrusion states: balanced alternation of protrusion types stabilizes migration trajectories, whereas clustering of similar protrusions leads to unstable trajectories and reduced directional persistence. Data are derived from 1,153 protrusions across 106 cells from independent experiments. Statistical comparisons were performed as described in Methods.

Across the control population, protrusion lifetime showed the strongest association with directional migration (Fig. 4B; Fig. S4A). All three temporal phases contributed positively to directional output, with retraction duration exhibiting the strongest phase-specific correlation. In contrast, retraction area was inversely associated with directional efficiency, indicating that prolonged and controlled retraction, rather than large collapse events, supports productive migration.

We next examined how NMII organization relates to protrusion persistence. Whereas instantaneous NMII levels at the front showed little association with protrusion lifetime, cumulative NMII engagement correlated strongly with protrusion persistence (Fig. 4C; Fig. S4B). Rear NMII load showed the strongest relationship with protrusion lifetime and retraction duration, indicating that sustained contractile engagement contributes to controlled protrusion retraction. These observations suggest that protrusion persistence reflects coordinated actomyosin activity across the cell rather than enrichment of NMII at a single pole. In this framework, front NMII contributes to protrusion stabilization, whereas sustained rear NMII engagement provides the strongest predictor of protrusion persistence and directional output.

Consistent with this interpretation, front NMII load was positively associated with directional output, whereas rear NMII load and front–rear polarity metrics showed stronger predictive power overall (Fig. 4D; Fig S4C). Thus, persistent migration emerges from coordinated engagement of spatially distinct actomyosin systems rather than from front accumulation of NMII alone.

To stratify protrusions based on directional behavior, we performed principal component analysis (PCA) on directional parameters and derived a composite directionality index (CDI) (Fig. S5A–D). Using a median CDI split, protrusions were classified into high- and low-directionality groups. High-directionality protrusions displayed significantly longer lifetimes driven primarily by extended retraction duration (Fig. 4E; Fig. S5E).

At the molecular level, these protrusions exhibited elevated rear NMII load and stronger rear-biased front–rear load differences (Fig. 4E). In contrast NMII lifetimes metrics showed weaker separation between high- and low-directionality protrusions (not shown).These results indicate that cumulative NMII engagement, rather than instantaneous intensity, best distinguishes protrusions that support directional migration.

We next asked how protrusions collectively generate persistent migration at the cell level. Cells were stratified into high- and low-directionality groups using the same PCA/CDI framework (Fig. S5F–I). Surprisingly, highly directional cells did not produce a greater fraction of directionality-supporting protrusions. In fact, such protrusions were proportionally more abundant in low-directionality cells (Fig. 4F), indicating that directional efficiency cannot be explained by simple enrichment of favorable protrusion types.

Instead, the key difference emerged in the temporal organization of protrusions. In highly directional cells, directionality-supporting protrusions occurred intermittently and were interspersed with other protrusion states. Consequently, the probability that a directionality-supporting protrusion was followed by another of the same type was significantly lower in highly directional cells (Fig. 4G). In contrast, low-directionality cells frequently produced consecutive runs of similar protrusion states and showed longer mean run lengths (Fig. 4H).

Together, these results indicate that persistent migration emerges from the temporal coordination of protrusion states rather than from the abundance of favorable protrusions. A conceptual model illustrates how balanced alternation of protrusion states stabilizes migration trajectories, whereas clustering of similar protrusions leads to unstable trajectories and reduced directional efficiency (Fig. 4I).

### Pharmacological perturbations reveal distinct temporal mechanisms controlling directional persistence

To probe the functional role of spatially organized NMII during migration, we perturbed NMII regulation using pharmacological inhibitors. Because NMII at the leading edge is phosphorylated on Ser19 (Fig. S2C), we inhibited Rho-associated kinase (ROCK) with Y27632^33,37^. The inhibitor was applied at a concentration that preserved motility in at least half of the cells (Fig. S6A), allowing analysis of partial perturbations rather than complete arrest of migration. As previously reported, ROCK inhibition markedly reduced NMII localization at the rear of migrating neutrophils (Fig. 5A; Movie S9). Quantitative analysis revealed a strong reduction in sustained rear NMII engagement, reflected by decreased rear NMII lifetime and a shift toward front-dominant NMII polarity (Fig. 5B and Fig. S6B).

**Figure 5.**
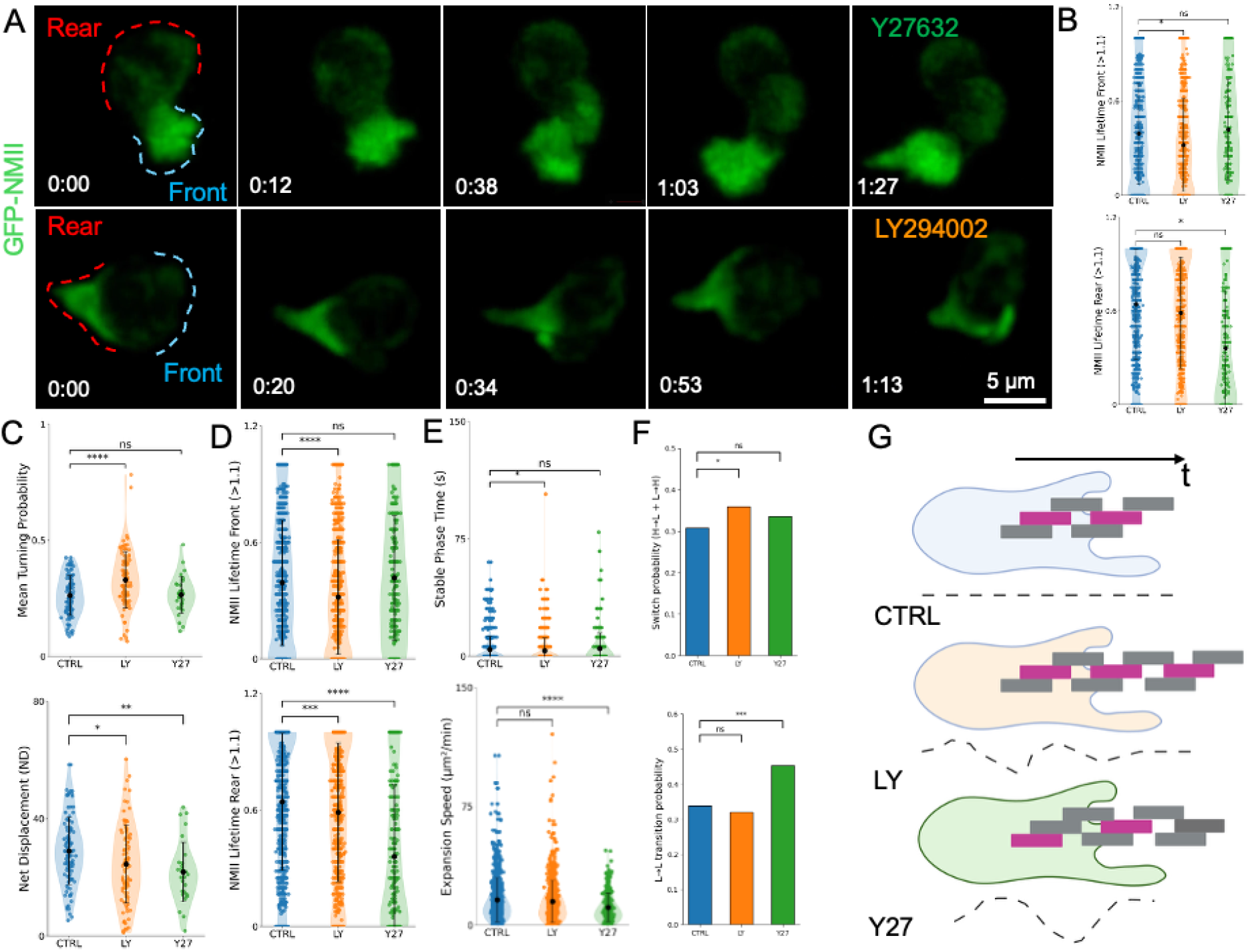
Pharmacological perturbations of NMII regulation reveal distinct mechanisms controlling directional persistence. (A) Representative time-lapse images of migrating neutrophils expressing GFP-NMII under inhibition of ROCK (Y27632) or PI3K (LY294002). Rear and front regions are indicated by red and blue dashed outlines, respectively.(B) Quantification of NMII persistence at the cell front and rear across conditions. Plots show the fraction of cells maintaining NMII levels above a high-threshold intensity (≥1.1) for more than 50% of the migration time, revealing reduced rear NMII persistence after ROCK inhibition and reduced front NMII persistence after PI3K inhibition.(C) Migration phenotypes measured at the cell level. ROCK inhibition significantly reduces net displacement (ND), whereas PI3K inhibition increases turning probability without strongly affecting displacement.(D) Protrusion-level NMII metrics showing that LY294002 reduces front NMII engagement during protrusion activity, whereas Y27632 reduces rear NMII persistence.(E) Structural protrusion parameters across pharmacological conditions. PI3K inhibition shortens the stabilization phase of protrusions, whereas ROCK inhibition reduces protrusion expansion speed. (F) Transition probabilities between protrusion states under different perturbations. Control cells show balanced transitions between high- and low-directionality protrusions, whereas PI3K inhibition increases switching and ROCK inhibition stabilizes low-directionality protrusion states.(G) Conceptual model illustrating how pharmacological perturbations alter the temporal organization of protrusion states. Balanced alternation of protrusion states supports persistent migration in control cells, whereas altered transition patterns under PI3K or ROCK inhibition destabilize directional persistence.Data represent individual cells or protrusions from independent experiments as described in Methods. Statistical significance was determined using the tests indicated in the figure.

We next inhibited PI3-kinase signaling using LY294002^38^, which regulates front polarity^39,40^. LY treatment reduced sustained NMII engagement at the leading edge without substantially altering rear NMII levels (Fig. 5A; Movie S10). Distribution analyses confirmed a reduction in front NMII lifetime across thresholds, consistent with destabilization of front-associated NMII engagement (Fig. 5B and Fig. S6B). Thus, the two inhibitors produced distinct spatial perturbations of NMII organization.

These spatial redistributions produced different migration phenotypes. ROCK inhibition significantly reduced net displacement (ND), indicating impaired forward propulsion, whereas PI3K inhibition had little effect on displacement but increased turning probability (Fig. 5C; Fig. S6C). These results indicate that rear NMII engagement primarily contributes to propulsion, whereas front NMII stability influences directional persistence. This reinforces the central discovery of the paper.

Consistent with these cell-level phenotypes, protrusion-scale measurements revealed distinct NMII perturbations. LY reduced front NMII engagement during protrusion activity, whereas Y27632 markedly reduced rear NMII persistence while leaving front levels largely unchanged (Fig. S6D). Importantly, protrusion lifetimes were largely unchanged across conditions (Fig. 5D), indicating that pharmacological perturbations primarily affected protrusion organization rather than intrinsic protrusion duration.

Instead, the inhibitors altered protrusion dynamics. PI3K inhibition shortened the stabilization phase of protrusions, whereas ROCK inhibition reduced protrusion expansion speed (Fig. 5E and Fig. S6D–F). Despite these kinetic effects, protrusion orientation remained unchanged (Fig. S6F), indicating that the perturbations primarily affected protrusion stability and propagation rather than directional bias.

Finally, we examined how these perturbations affected the temporal organization of protrusion states. Under control conditions, protrusion states alternated in a balanced manner, consistent with the intermittent protrusion sequences observed in highly directional cells. LY treatment increased switching between protrusion states, whereas ROCK inhibition stabilized runs of low-directionality protrusions (Fig. 5F). These results indicate that directional persistence is controlled not by protrusion lifetime alone but by the temporal sequencing of protrusion states. Together, these experiments demonstrate that perturbing NMII regulation produces spatially distinct actomyosin phenotypes that propagate from protrusion dynamics to whole-cell migration. PI3K inhibition destabilizes front NMII engagement and increases protrusion-state switching, whereas ROCK inhibition disrupts rear contractile persistence and reduces forward propulsion. These findings support a model in which directional persistence emerges from the coordinated temporal organization of protrusion states governed by spatially distinct NMII pools.

### Myosin II forms lattice-like assemblies at the leading edge of migrating neutrophils

Because our analyses indicated that elevated NMII levels at the leading edge are associated with larger protrusions, we next examined how NMII is structurally organized at this location. To address this question, we used high-resolution spinning-disk microscopy and structural illumination microscopy (SIM) to visualize NMII organization at the plasma membrane of migrating neutrophils.

At the leading edge of cells migrating in three-dimensional collagen matrices, NMII formed discrete assemblies composed of nodes connected by short filamentous structures (Fig. 6A). Similar structures were observed in vivo (Fig. 6B and Fig. S7A). These assemblies appeared as triskelion-like arrangements of NMII mini-filaments radiating from central nodes, resembling lattice-like NMII organizations previously described on large secretory granules^41^.

**Figure 6.**
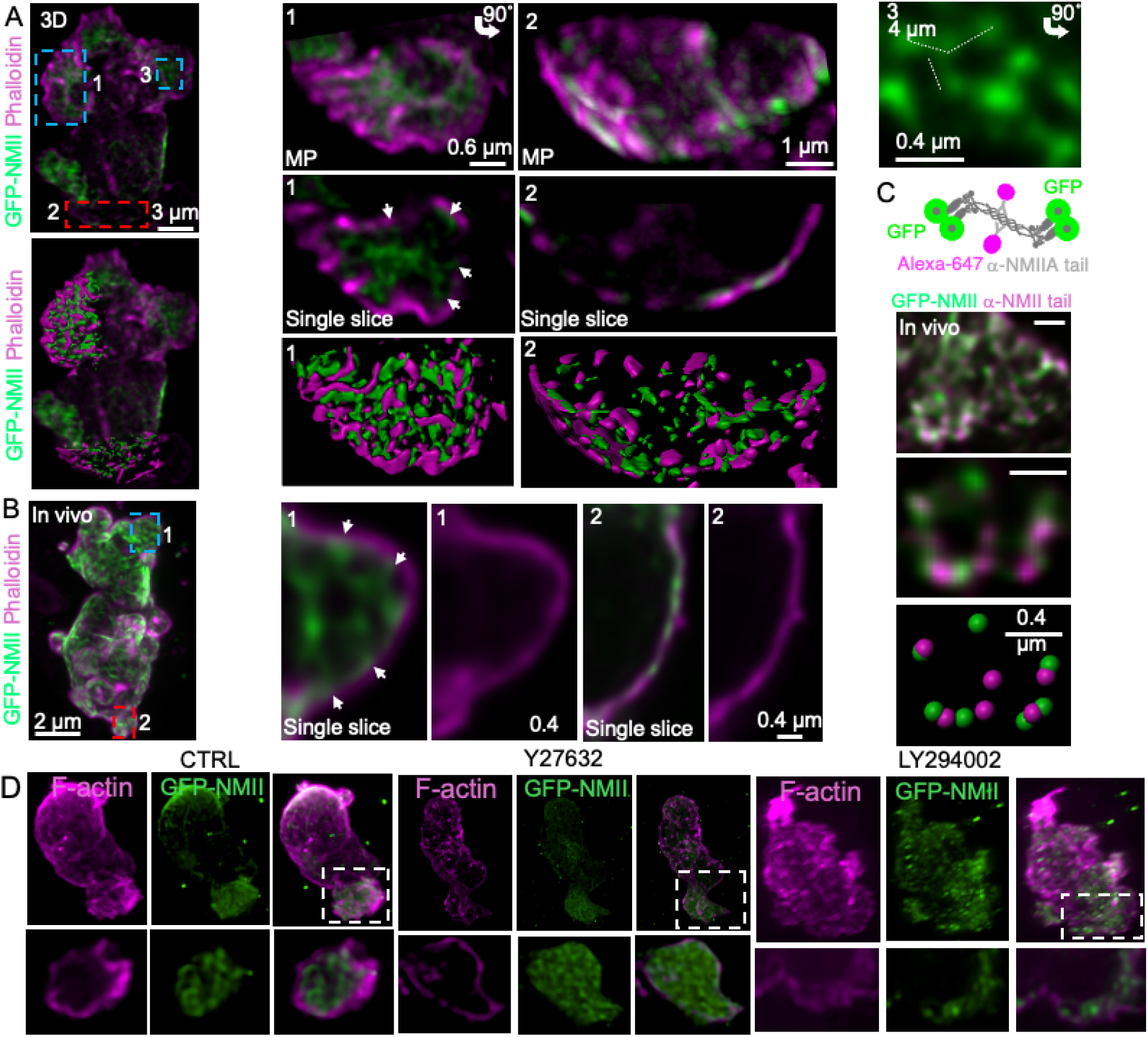
Myosin II forms lattice-like assemblies at the leading edge of migrating neutrophils. (A) Organization of NMII in neutrophils migrating within three-dimensional collagen matrices. Maximum-intensity projections of fixed GFP-NMIIA neutrophils labeled with phalloidin (magenta) show NMII distribution at the leading edge and rear of the cell. Insets show higher-magnification views of the leading edge (1) and rear (2). Upper panels display maximum projections, middle panels show single optical slices highlighting NMII puncta (arrows), and lower panels show isosurface renderings illustrating the organization of NMII assemblies relative to cortical actin. (B) Organization of NMII during neutrophil migration in vivo. Single optical slices of representative cells show discrete NMII assemblies at the leading edge and cortical NMII bundles at the rear. (C) Visualization of NMII mini-filament organization using antibody labeling of the NMIIA tail domain together with GFP-NMIIA, revealing triskelion-like arrangements consistent with interconnected NMII mini-filaments. (D) Effects of pharmacological perturbations on NMII organization. Representative images of neutrophils treated with ROCK inhibitor (Y27632) or PI3K inhibitor (LY294002) and labeled for NMII (green) and F-actin (magenta). PI3K inhibition disrupts the lattice-like organization of NMII at the leading edge, whereas ROCK inhibition increases NMII density at the front and reduces cortical NMII at the rear. Scale bars as indicated.

Visualization of NMII motor heads and tail domains further supported this organization. In many cases, NMII heads partially overlapped with cortical F-actin, suggesting that these assemblies represent sites of interaction between the actomyosin cortex and the plasma membrane (Fig. 6A,B). In contrast, NMII at the rear of migrating cells displayed a markedly different organization, forming bundles of parallel mini-filaments aligned along the cortex and closely associated with F-actin (Fig. 6A and Fig. S7B–C), consistent with their contractile function during rear retraction.

Perturbations that altered NMII polarity also modified the organization of these assemblies. PI3K inhibition with LY294002 disrupted the lattice-like architecture at the leading edge while leaving cortical NMII largely intact (Fig. 6D). Conversely, ROCK inhibition with Y27632 produced a denser NMII network at the front while reducing cortical NMII at the rear (Fig. 6D). Importantly, neither perturbation altered the global organization of cortical F-actin (Fig. S7D–E), indicating that the observed effects primarily reflect changes in NMII organization rather than secondary disruption of the actin cytoskeleton.

Together, these observations indicate that NMII adopts two distinct supramolecular organizations during neutrophil migration. At the leading edge, NMII forms lattice-like assemblies associated with protrusive regions of the cell cortex, whereas at the rear it forms contractile bundles aligned with cortical actin. These spatially distinct architectures suggest that front and rear NMII systems contribute differently to the mechanical coordination of protrusion stabilization and cell propulsion during migration (Fig. 6D).

## Discussion

Directed neutrophil migration in tissues is a central component of innate immune responses, yet the mechanisms that stabilize persistent movement in complex three-dimensional environments remain incompletely understood. Most current models of neutrophil motility derive from two-dimensional systems, where non-muscle myosin II (NMII) is primarily viewed as a rear-localized contractile motor responsible for uropod retraction and forward translocation^15^. Here, by combining intravital subcellular microscopy with quantitative analysis of membrane and cytoskeletal dynamics, we identify a previously unrecognized spatial organization of NMII during neutrophil migration in vivo and in three-dimensional matrices. Our results indicate that NMII operates through spatially distinct architectures at the front and rear of migrating cells that coordinate protrusion dynamics and directional persistence.

A central finding of this study is that NMII adopts two distinct supramolecular organizations during neutrophil migration in three-dimensional environments. At the leading edge, NMII assembles into lattice-like structures composed of interconnected mini-filaments, whereas at the rear it forms parallel cortical bundles associated with contractile actin networks. These two architectures differ not only structurally but also functionally. Unlike the well-characterized rear bundles that generate contractile forces, front-localized NMII forms triskelia-like assemblies of bipolar filaments that appear associated with protrusive regions of the cell cortex. This structural organization correlates with increased protrusion expansion area and prolonged protrusion growth phases, without directly triggering membrane retraction. These observations suggest that NMII at the leading edge primarily contributes to the mechanical stabilization of expanding protrusions rather than to contractile collapse of the cortex.

Such stabilization is likely to be particularly important in three-dimensional environments, where cells migrate through irregular pore spaces and heterogeneous extracellular matrices. In these contexts, protrusions must persist long enough to engage the surrounding matrix and guide cell movement. By contrast, in two-dimensional migration systems actin-driven lamellipodia dominate protrusion dynamics and NMII remains largely confined to the rear of the cell^15,16,35^. The presence of NMII assemblies at the front of migrating neutrophils therefore reflects a fundamental difference between migration in simplified planar environments and migration within complex tissue-like matrices.

Our quantitative analyses further reveal that protrusion persistence is a major structural determinant of directional migration. Protrusion lifetime shows a strong association with directional efficiency, and sustained NMII engagement —particularly at the rear of the cell— correlates with prolonged protrusion retraction phases. However, persistent migration cannot be explained simply by the presence of favorable protrusions. Instead, highly directional cells display a characteristic temporal pattern in which different protrusion states alternate over time. In contrast, cells with lower directional efficiency tend to produce extended runs of similar protrusion states, resulting in unstable trajectories and increased turning behavior. These findings indicate that directional persistence emerges from the temporal organization of protrusion dynamics rather than from the properties of individual protrusions alone.

In this framework, the spatial organization of NMII provides a mechanism for coordinating protrusion behavior across the cell. The lattice-like assemblies at the leading edge appear to stabilize protrusive structures and reduce stochastic fluctuations in front dynamics, whereas rear NMII bundles generate sustained contractile forces that support retraction and forward propulsion. Directional migration therefore arises from the coordinated activity of these spatially distinct actomyosin systems. Rather than acting independently, the front and rear NMII pools appear to influence protrusion dynamics through complementary mechanical roles that stabilize migration trajectories. Pharmacological perturbations further support this interpretation by revealing distinct regulatory pathways controlling these NMII populations. Inhibition of PI3-kinase selectively destabilizes front-associated NMII engagement and disrupts lattice organization, while largely sparing rear contractile NMII. Conversely, inhibition of ROCK primarily affects the rear contractile pool and increases lattice density at the leading edge. These perturbations produce distinct migration phenotypes: PI3K inhibition increases turning behavior without substantially reducing displacement, whereas ROCK inhibition impairs forward propulsion. Importantly, protrusion lifetimes remain largely unchanged across perturbations, indicating that these signaling pathways primarily influence the temporal organization of protrusion states rather than the intrinsic duration of protrusive events.

These findings place NMII downstream of established polarity pathways that regulate neutrophil migration. PI3-kinase signaling has long been implicated in leading-edge polarity and directional sensing^39,40^, typically through control of actin polymerization and membrane lipid gradients. Our results extend this framework by suggesting that PI3K-dependent pathways regulate NMII organization at the leading edge, thereby stabilizing protrusions that align with the migration axis. Conversely, ROCK-dependent signaling appears to sustain the rear contractile system that drives forward propulsion. The coordinated regulation of these two NMII architectures therefore provides a mechanism by which cells integrate polarity signaling with mechanical control of migration.

Although neutrophils express only the NMIIA isoform, the behavior of front-localized NMIIA observed here shares conceptual similarities with functions attributed to NMIIB in other cell types. NMIIB has frequently been associated with maintaining polarity and sustaining tension over longer timescales, whereas NMIIA is typically linked to rapid contractile dynamics^42,43^. The lattice organization and prolonged residence time of NMIIA at the leading edge suggest that supramolecular assembly may enable a single NMII isoform to perform multiple mechanical roles depending on spatial context. This highlights the importance of higher-order cytoskeletal organization in determining myosin function.

Interpretation of the pharmacological perturbations presented here should nevertheless be considered within the limitations of inhibitor-based approaches. Although PI3K and ROCK inhibition produced reproducible and spatially distinct phenotypes, pharmacological perturbations can influence multiple downstream pathways and may not fully isolate specific regulatory mechanisms. Additional inhibitors targeting NMII regulation or upstream signaling components were evaluated but produced variable or inconclusive effects and were therefore not pursued further. The perturbations presented here should therefore be viewed primarily as tools that reveal the functional contributions of spatially organized NMII systems rather than as definitive pathway-specific manipulations.

In conclusion, our results reveal that neutrophil migration in complex environments relies on spatially distinct NMII architectures that coordinate protrusion dynamics across time. By linking cytoskeletal organization to the temporal sequencing of protrusion states, this work identifies a previously unrecognized mechanism that stabilizes directional migration during immune cell motility in tissues. More broadly, these findings suggest that regulation of protrusion persistence—rather than protrusion initiation or retraction alone—may represent a fundamental principle controlling directional migration in physiologically relevant environments.

## Acknowledgments

This work was supported in part by the Partnerships Program in the Office of Intramural Training & Education of the National Institutes of Health. R.W. was supported by the National Institutes of Health, National Cancer Institute, Center for Cancer Research Intramural Research Program (ZIA BC 011682). We thank Michael Kruhlak, Langston Lim, and Dr. Andy Tran at the CCR Confocal Microscopy Core and Sandep Pallikkuth and Itoro Akpan at the LCMB Microscopy Core for their assistance with light microscopy. The content of this publication does not necessarily reflect the views or policies of the Department of Health and Human Services, nor does mention of trade names, commercial products, or organizations imply endorsement by the U.S. Government. All schematics and cartoons were created using BioRender.

## Author Contributions

N.M. and R.W. designed the experiments. N.M., E.C., T.M., Y.N., B.S. and W.W. performed the experiments. C.P. and W.L. provided intellectual insights throughout the project. N.M., D.C., L.W., and R.W. analyzed the experiments. N.M., D.C., and R.W. wrote the manuscript. All authors read and approved the final manuscript.

## Material and methods

### Experimental Models Mice

All experiments were approved by the National Cancer Institute (National Institutes of Health, Bethesda, MD, USA) Animal Care and Use Committee and were compliant with all relevant ethical regulations regarding animal research. FVB, C57Bl6/NJ, mTmG and cGFP were in-house produced from commercially available mice strains from Jackson Laboratory. *GFP-NMIIA knock-in* mice were gifted by Robert Adelstein (NHLBI, National Institutes of Health). Mice enrolled in experiments were between 2-6 months of age, weighted 20-40 g and both genders were used without any gender specific effects observed.

### Reagents

Antibodies to P-ERK (#4370), total-ERK (#4695), P-MLC2 Ser19 (#3671 & #3675), total-MLC2 (#8505) were from Cell signaling Technology, Ct-NMIIA antibody was from Biolegend (PRB-440P). Recombinant fNLFNYK and WKYMVm were from Santa Cruz Biotechnology and Abcam respectively. Y27632, Ly294002 and ML-7 were from MedChem Express, Nitro-Blebbistatin was from Cayman, EHT186 was from R&D systems. DAPI,, Celltracker-Green and Alexa conjugated phalloidin and secondary antibodies were from Invitrogen. Atto647 conjugated Phalloidin was from Sigma-Aldrich. Cellmask-Deep Red was from Cytoskeleton.

### Primary mouse and human neutrophil isolation

Mouse neutrophils were purified from mouse front and hind limbs bone marrow. Briefly, bone marrow cells were flushed out with a 27GA syringe using HBSS without Red Phenol, Calcium and Magnesium (Invitrogen), supplemented with 10mM HEPES (Quality Biological). Cells were resuspended in ACK buffer (Gibco) to lyse Red Blood cells (RBC) then washed with HBSS before to be layered on a discontinuous step gradient formed with 1119 and 1077 Histopaque (Sigma-Aldrich). After 30min centrifugation at 1000g, primary neutrophils were collected at the 1119 and 1077 interface, washed 2 times and then, resuspended in HBSS to be counted.

Human neutrophils (PMNs) were obtained from heparinized whole blood of healthy human donors as part of the NIH Blood Bank research program. Whole blood was centrifugated on Histopaque 1077 and the fraction containing RBC and PMNs was subsequently sedimented on a 3% Dextran solution (0.5 MDa, Sigma-Aldrich). RBC were lysed by multiple cycles of hypotonic-isotonic conditions (1 volume of 0.2% NaCl for 30 s neutralized by 1 volume of 1.6% NaCl) and centrifugations.

All purified neutrophils were kept on a rotating rotor at RT in HBSS where they were labeled and/or treated with inhibitors. In vitro assays were performed at 37°C unless specified.

### Imaging

#### Microscopes setups and image acquisition

##### Multiphoton microscope

Inverted laser-scanning MPE-RS Olympus microsocope equipped with a Spectra Physics Insight DS+ tunable laser and a heated stage/objective setup (Okolab and Biotechs respectively). Excitation was performed at 900 nm and emitted light was collected by an appropriate set of mirrors and filters on GaAs detectors (bandpass filters: Blue = 410–460 nm, Green = 495–540 nm, FarRed = 670-730 nm). Objectives used for imaging were Olympus USPLAPO 30x (NA 1.05) and 40x (NA 1.25) silicone oil immersion for respectively low and high magnification. Imaging was performed using resonant scanner for live imaging and galvo scanner for still imaging acquired volumes. Images were acquired using the Olympus Fluoview software.

##### Spinning Disk

Inverted Nikon Eclipse Ti2 microscope equipped with Yokogawa CSU-X1 spinning-disk confocal system and a Hamamatsu FusionBT camera. Objectives used for the imaging were SR HP Plan Apo Lambda 100XC Silicone oil (NA 1.35) or 60X Oil immersion (NA 1.4). Live imaging was performed at 100-300ms exposure time per frame, with Z-stacks acquired at 0.1-0.5 µm apart and still imaging at 100-300ms exposure time per frame with Z-stacks acquired at 0.05-0.3 µm apart. Images were acquired using the Nikon Elements software and when mentioned, imaged were processed post-acquisition in the Elements software to denoise the image using the Nikon Denoise.ai function and was followed by 3D deconvolution using the Richardson– Lucy algorithm with (1) an additional step for pre-noise estimation and (2) automatic determination of the number of iterations.

Both spinning disk and 2-photon microscopes were equipped with fine controlled heating system of the stage and of the objective to ensure a stable 37°C imaging temperature for live experiments.

###### Other microscopy setups

Deep-SIM: Inverted Nikon Eclipse Ti2 microscope equipped with a super-resolution module DeepSIM (CrestOptics), a Hamamatsu Fusion BT camera and equipped with a 100x SR HP Plan Apo Lambda Silicone oil immersion objective (NA 1.35).

##### Sora

Inverted Nikon Ti2 CSU-W1 microscope equipped with a Yokogawa SoRa CSU-W1 spinning disk unit, a Photo-metrics BSI sCMOS camera and a 60x SR Plan Apo immersion objective (NA 1.27).

Images acquired with those systems were further viewed and rendered for publication using Imaris (Bitplane) or Fiji (ImageJ) or analyzed using MATLAB (Math Works) as described further.

#### Intravital / ISMic imaging

Neutrophil injection in the ear of a recipient mouse were performed as described previously^19^. Briefly: albino recipient mouse was lightly anesthetized by isoflurane inhalation (Forane; Baxter Healthcare Corp.) and received 3 to 5 injections of a neutrophil suspension (2-4 µl, approx 2-3.10^5^ cells per injection) in a shaved ear with a 33 G needle to reduce tissue lesion by the injection. After a recovery period, recipient mouse was anesthetized with a Ketamine/Zylazine/Acepromazine solution (80 mg.kg^-1^, 2 mg.kg^-1^ and 4 mg.kg^-1^ respectively, VET One) and installed on a custom designed holder to reduce motions. Imaging was performed with the multiphoton microscope setup for either live (10-20 z-steps, z-spacing = 1 µm, every 5-10 s) or still imaging of tissue microenvironment and collagen organization (z-spacing = 2 µm).

During the experiment, any sign of bleeding, swelling or inflammation of the ear were a direct parameter of exclusion from the experiment.

#### Under agarose assays

Briefly, #1.5 glass bottomed 8 wells chambers were coated over-night with type I collagen (Rat tail collagen, Corning) and 5% BSA (Sigma). A 1% agarose gel was casted in each well and the chamber was kept at 4°C to allow gelling. Once solid, gels were pierced by two 1mm diameters holes, separated by 2.5mm and gels were then equilibrated at 37°C for 30min before the seeding in one of the well. After 20 min, the second well was filled out with a 0.3 µM WKYMVm solution and live imaging was performed using previously described Olympus two-photon microscope equipped with a 30x silicon oil immersion objective.

In some instances, samples were fixed using a 4% paraformaldehyde solution (Electron Microscopy Sciences) and ultimately used for immunofluorescence experiments.

#### 3D collagen gels assays

Embedded neutrophils in 1.7mg/ml collagen-I hydrogels were used to recreate complex 3D environments. Rat tail type-I collagen (Corning) was used to create 3D hydrogels either on a glass bottom dish for live imaging or in an Eppendorf for biochemical analysis. Cells were resuspended to the desired concentration in a HBSS solution containing 20mM HEPES at pH 7.4 and in parallel, collagen was diluted to a 2x final concentration in HBSS. To ensure optimum collagen polymerization and cell viability, cell suspension and collagen mix were kept at 4°C and mixed at the last minute in presence of a sufficient volume of 0.1N NaOH to bring the final pH of the solution to 7.4. Gels were allowed to polymerize for 30 min at 37°C and cells were then exposed to inhibitors and/or chemoattractant. Live imaging was performed with the inverted two-photon microscope or, for higher temporal and spatial resolution, with the spinning disk microscope. In some instances, samples were fixed using a 4% paraformaldehyde solution (Electron Microscopy Sciences) and ultimately used for immunofluorescence experiments.

### Sample Preparation

#### Cardiac fixation, OCT embedding and cryosectioning

To perform immunofluorescence on tissues, ears are collected after cardiac perfusion of fixative: left ventricule of the heart is punctured and 10 ml of saline solution is injected to wash out the blood from the right atrium, followed by 10 ml of fixative (4% paraformaldehyde in PBS buffered at pH 7.3). Ears are then cut off and placed in fixative solution for 1 hour at room temperature then washed 3 times. After fixation, ears were put through a sucrose gradient (10% → 20% → 30%) for cryoprotection. The tissue was then placed in a mould containing OCT and allowed to freeze on dry ice.

To obtain higher imaging resolution, 3D gels, after paraformaldehyde fixation and rinses were also embedded in OCT for cryosection (to improve optical resolution compared to whole mount).

Cryosection was performed on a Leica cryostat microtome (CM1860, −13°C, 15 µm thickness) and sections were collected on a charged slide and then processed for immunofluorescence.

#### Immunofluorescence

After fixation, samples were quickly washed 3-5 times with 0.2M HEPES pH 7.2 to remove any traces of fixatives and either whole mounted or frozen in OCT for cryosection. Either whole samples or cryosections were then permeabilized for 30min (0.5% Triton X-100 and 0.5% Saponin solution in 0.2 M HEPES) and blocked in 10% NGS for 30min at room temperature. Samples were then incubated with primary antibody for 48h at 4°C (whole mount) or 1h at room temperature (cryosections), rinsed 3 times and stained with appropriate secondary antibody for 1h at RT (Anti-Rabbit IgG and anti-Mouse IgG, 1/500 dilution, ThermoFisher) and counterstained with Phalloidin (1/100 dilution, Sigma) and/or DAPI. Primary antibody used were the following: MLC Ser19 (rabbit and mouse, CST, 1/100 dilution), Ct-NMIIA (rabbit, Biolegends, 1/300 dilution). Whole mount and cryosections were imaged using the spinning disk except if specified otherwise.

#### Western blotting

Samples were directly lysed in a Laemmli solution containg 5x protease and phosphatase inhibitors (cOmplete and PhosSTOP respectively, Roche). Lysates were snap frozen in liquid nitrogen, sonicated and centrifuged at 20,000 g for 15 min at 4°C. Samples were then loaded on a 4-20% SDS-PAGE gel and transferred on a PVDF membrane. Membrane was then blocked for 1 hour at room temp with 5% BSA in TBST and incubated overnight at 4°C with the corresponding primary antibody directed against total ERK, P-ERK, total MLC and MLC-Ser19 (CST). Blots were incubated with horseradish peroxidase–conjugated secondary antibodies (CST) for 1 h at room temperature and signal was reavealed using an ECL reagent (SuperSignal™ West Pico PLUS, Thermo) on a Chemidoc (BioRad)

#### Imaging analysis and quantitative analysis

#### Four Dimensional (4D) Image processing and quantification

##### Cell segmentation and tracking

Representation of cell segmentation: briefly, after cell segmentation, the 2D cell boundary was divided in 100 equal segments to finally define 100 sections within the 2 µm subcortical area. Those sections and the associated voxels that can be z-projected in these sections were then distributed as front (30 sections), rear (30 sections) and 2 sides (20 sections each) according to the cell direction migration direction, as represented on the cartoon. Remaining of the cells is considered as uncharacterized.

4D images were imported to MATLAB with the Bio-Formats package. Further analyses were accomplished with customized MATLAB codes. For the videos acquired in vivo, a correlation-based frame registration and motion correction was applied to the collagen I-SHG channel. For all videos, at each frame, the segmentation of the 3D objects in the GFP-NMIIA channel was achieved with the Otsu algorithm. The identified objects in the video were tracked with the minimum-displacement criterion applied to their geometrical centroids from each frame to the next frame. 2D boundaries of the max-projections of the objects were extracted. Snake algorithm was applied to improve the precision of the boundary to the subpixel level and define 100 consecutive points along the boundary with even spacing. The object boundaries and the tracking identification indices were overlayed with the original 4D images and output for visualization and manual inspection. In the manual inspection, tracked objects were sorted as active individual cells or non-activated round and still cells (Fig. S5). Non-cell objects, touching objects, and objects with defects in the boundary identification were excluded.

The migration path of each individual migrating cell was determined by the trace of the geometric centroid frame to frame. The average cell migration speed overtime of each cell was defined as *l*/Δ*T*, where *l* is the length of the entire migration path and Δ *T* is the time deference of the start and end of the cell migration path. Within the active cell subpopulation, only motile individual cells with no less than 10 frames and with average speed no less than 0.1 μm/s were further analyzed. Active cells with average speed less than 0.1 μm/s were categorized as jiggling cells. The fractions of the still, jiggling, and motile cells were calculated by the number cells in each subtype over the total number of tracked cells. The instant cell migration speed was measured as the length of the framewise displacement divided by the frame rate. The overall cell migration tortuosity was defined as *l*/|*s*|, where *s* is the displacement vector from the start to the end of the cell migration path. The migration direction *n* at each time point was calculated as the direction of the displacement in the next two frames to smooth the effect of subcellular dynamics on the identification of entire cell migration direction. To measure the instant deviation of the migration direction, at each time point the deviation angle of the migration direction was defined as the angle between *n* and *n*_*next*_, where *n*_*next*_ is the migration direction calculated for the next time point. The cell migration direction persistence with time offset Δ*t* was measured by the auto-correlation coefficient of migration direction as:

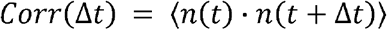

Here ⟨ ⟩ denotes the averaging operation at all time points *t* if Δ*t* ≥ 0 over all cells in the population (Fig. 3B).

#### Quantification of subcellular dynamics during cell migration

The subcellular dynamics of cell shape during cell migration were calculated by tracking the motions of the 100 boundary points from one frame to the next with the minimum displacement criterion. For each boundary point at a certain time point, if the local cell boundary moved outward from the cell, a positive sign was assigned to the boundary point motion to characterize the local protrusion. Vice versa, a negative sign was assigned to the local retraction if the local cell boundary moved inward to the cell. The local area change at each boundary point at a certain time point was defined as the area of the polygon of which vortices are the tracked boundary points of interest in the current and next frame, the middle points between the points of interest and their neighbor boundary points on the clockwise side, and the middle points on the counterclockwise side. The same positive or negative signs of the motion of the center boundary point were given to characterize the local protrusion or retraction. The local membrane curvature at each boundary point for each frame was measured by fitting its neighbor boundary points to a circle (5 per side; total 11) with the Pratt circle fitting method.

To measure the local cortical GFP-NMIIA distribution of the cell at each time point, the depth of the cell cortex was set as 2 μm^44^ so that the inner boundary of the cell cortex was identified within the 2D boundary of the cell max-projection. 100 inner boundary points were then identified as the points on the inner cortex boundary closest to the 100 cell boundary points. With the boundary points on the inner cortex boundary and the cell boundary, 100 local cortical sections along the cell cortex were segmented (Fig. 1H). At each time point, for the *k*th cortical section, we calculated the average raw NMIIA-GFP local cortical intensity *I*_*raw,k*_(t) of *N*_*k*_ (*t*) 3D cell voxels that can be z-projected in that section, and the average raw GFP-NMIIA intensity of all voxels *I*_*raw,mean*_(*t*) at time *t* in the cell. For each cell, we identified the minimum level of the local raw cortical NMIIA-GFP intensity *I*_*raw,min*_ = min_*k,t*_ *I*_*raw,k*_(*t*) over all cortical sections at all time points. The normalized local cortical NMIIA-GFP level in the *k*th cortical section at time t was defined as:

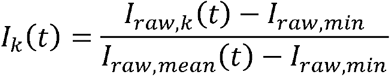

To smooth the noise caused by the limited number of voxels in each cortical section, for the *k*th boundary point the local cortical GFP-NMIIA level was calculated as the average of the two cortical sections on its clockwise side and the two cortical sections on its counterclockwise side as:

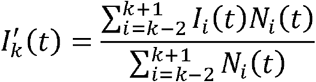

In the 100 boundary points, the 30 consecutive points facing the historical trace of cell migration path (e.g. the opposite of the entire cell migration direction) were identified as the rear of the cell boundary. The associated 30 cell cortical sections were identified as the rear cortical part of the cell. The 30 consecutive boundary points and associated cortical sections on the opposite side of the rear were identified as the front of the cell. Between the front and the rear of the cell, the 20 boundary points and associated cortical sections on each side were identified as the two sides of the cell. The widths of the front or rear of the cell were calculated respectively as the span distances of the front or rear boundary points on the direction perpendicular to the cell migration direction. The normalized cortical GFP-NMIIA level at the certain part of the cell cortex at each time point was calculated as:

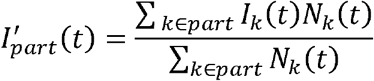

where the part refers to either front, rear, or side cortical regions of the cell.

We also calculated the average raw GFP-NMIIA local intensity *I*_*raw,unassigned*_ (*t*) for the 3D region unassigned to neither front, rear, or side cortical regions comprised of *N*_*unassigned*_ (*t*) voxels. The normalized GFP-NMIIA level in the unassigned part of the cell was quantified as:

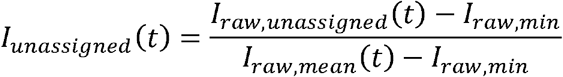

#### Quantification of correlations of subcellular dynamics

To quantify how cell front protrusions and rear retractions are coordinated during cell migration, we followed the methods described by Tsai et. al^15^. The Pearson correlation coefficient of any two time-dependent measurements *x*(*t*) and *y*(*t*) of a cell with time offset Δ*t* is defined as:

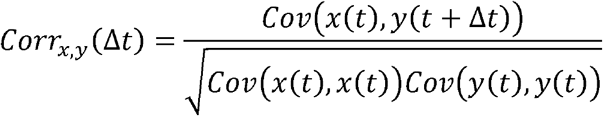

where *Cov*(*x,y*) is the covariance of variables *x* and *y*. For each cell, the Pearson correlation coefficient of the magnitudes of the total front protrusion activities and rear retraction activities was calculated with respect to a varying time offset. The bootstrap analyses were also applied to obtain the 95% confidence intervals at every Δ*t*. The time offset of the peak correlation was taken as the characteristic time difference of the onsets of front protrusion and rear retraction activities if the lower limit of the 95% confidence interval of the peak correlation is higher than 0.2. This threshold was taken to assure the significance of the correlation. A positive time offset of the peak correlation suggests the rear retraction lags the front protrusions and vice versa. The characteristic time offset of the front protrusion and rear cortical GFP-NMIIA level fluctuation was calculated in the similar way for each cell. The distributions of the time offsets were calculated to investigate whether the front protrusion activities lead the rear retraction activities or the rear cortical GFP-NMIIA level fluctuations on the populational level.

The motion direction of the cell front at each frame *n*_*front*_(*t*) was defined as the motion direction of the geometrical centroid of the polygons encompassed by the front boundary points. The motion direction of the cell rear at each frame *n*_*rear*_ (*t*) was defined in the similar way. The correlation of the front and rear motion directions during the cell migration was calculated as:

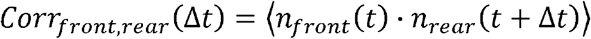

Here ⟨ ⟩ denotes the averaging operation over all cells in the population at all time points. Both Δ*t* ≥ 0 and Δ*t* < 0 were quantified to investigate whether the cell front steers the motion of the cell rear.

We also quantified the coordination of the local GFP-NMIIA cortical distribution and local membrane remodeling dynamics. For each migrating cell, the Pearson correlation coefficients of the local cortical NMIIA-GFP levels and the local boundary curvatures with time offset Δt = 0 were calculated over all time points and boundary points within the entire cell boundary, the front, the side, or the rear respectively. To specifically identify whether the GFP-NMIIA cortical distribution is correlated with positive or negative local curvature, the covariance calculation was modified so that the local curvature was centralized at 0 instead of the mean.

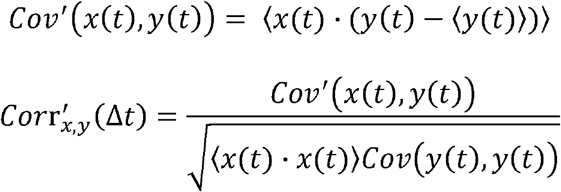

Here ⟨ ⟩ denotes the averaging operation over all cells in the population at all time points and associated boundary points in different regions. *x*(*t*) is the local curvature and *y*(*t*) is the local GFP-NMIIA level at a certain time point in this case. The correlations of GFP-NMIIA localization and local cell membrane motion of different cell cortical regions of interest were quantified with the Pearson correlation coefficient in the similar way where the local cell membrane motion was centralized at 0 instead of the mean as well.

#### Representation of single cell measurements with temporal fluctuations

Simply averaging over time neglects the temporal fluctuations of several single cell measurements such as the instant cell migration speed and GFP-NMIIA levels. To represent the level of a certain single cell measurement *x* with temporal fluctuations, we quantified the time fractions *f*_*x*_(*x*_*thr*_) of the cell to maintain *x* > *x*_*thr*_, where threshold *x*_*thr*_ was varied within the full meaningful range of the measurement to obtain objective quantifications. For the cell population of each independent experiment, we quantified the fraction of cells *F*_*x*_ (*x*_*thr*_,*t*_*thr*_) with time fraction *f*_*x*_ (*x*_*thr*_) > *t*_*thr*_. In this manuscript, we picked *t*_*thr* =_ 0.5 (or *t*_*thr*_ 0.67 in Fig. S2C), and plotted the average cell fraction over different independent experiments 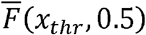 as a function of the varying *x*_*thr*_ with standard error of the mean (SEM). For different cell populations in comparisons, 1-way ANOVA tests were applied to *F*_*x*_(*x*_*thr*_, *t*_*thr*_) at every *x*_*thr*_ sampling points over independent experiments to investigate whether the differences between populations are significant. In this way, we objectively manifested the difference between cell populations taking both single cell variability and temporal fluctuation into consideration, without any need of *a priori* or arbitrary justifications about the characteristic level of the associated measurement.

#### Quantification of dynamics of the high-curvature protrusions

A newly generated local high-curvature protrusions on the cell boundary was identified with the following criteria: first it is comprised by consecutive cell boundary points, of each curvature higher than 0.2 μm^-1^, and second the center of this region doesn’t overlap with any region satisfying the first criterion in the previous frame according to the boundary tracking results. The center of each high-curvature protrusion region was defined as the boundary points of the highest curvature with a half width of 2 boundary points on each side. The frequency of the new protrusions of each cell was calculated as the total number of new protrusions divided by the total time of cell migration. The protrusion and retraction rates of each tracked protrusion at each time point were characterized as the total local area change of the protrusion center. The lifetime of the high-curvature protrusion was calculated as the time taken by the center of the high-curvature region to keep protruding or staying before retraction. The tracked protrusions were classified as stable protrusion with lifetime > 6s (1 frame) and nonstable transient protrusions with lifetime < 6s (1 frame). Within the subpopulation of the stable protrusions, we also identified long-life protrusions with lifetime > 30s. The frequencies of the mentioned subtypes of protrusions were calculated respectively as the total number of the new protrusions of each type divided by the total cell migration time. For each stable protrusion, the expansion rate of the protrusion was defined as the average protrusion rate prior to the steady stage at which time point the protrusion rate started to be less than. The disassembly rate was calculated as the average area change of the protrusion center after the protrusion lifetime finished. At each time point during the protrusion lifetime, The GFP-NMIIA level of the high curvature protrusion was calculated as the average normalized local cortical NMIIA-GFP level within the center of the protrusion as:

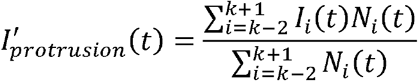

Where *i* is the index of the boundary point with the highest curvature of the protrusion. The mean GFP-NMIIA level of the protrusion was the mean of 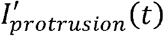 during the protrusion lifetime. The GFP-NMIIA localization lifetime was calculated as the longest consecutive period of time in which the GFP-NMIIA level 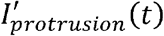 is always higher than threshold 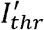 during the protrusion lifetime.

Data were visualized and plotted with Prism (GraphPad) except heat maps rendered using MATLAB. Statistical analysis tests were performed, and specific tests were picked as appropriated for each figure.

#### Global alignment of protrusion windows

All protrusion-resolved analyses were performed using global frame alignment relative to the full cell trajectory rather than sequential indexing from the start of each protrusion window. Each migrating cell was imaged over a global time series of frames, where the first frame corresponds to the initial coordinate of the tracked cell trajectory.

For each protrusion, the onset frame was defined using the “Frame” column corresponding to the global coordinate series. The protrusion start index was therefore defined as:

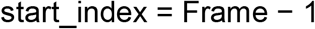

where indexing was converted to zero-based coordinates for computational analysis. Protrusion duration was determined from the measured protrusion lifetime and converted to frame units using the imaging frame rate:

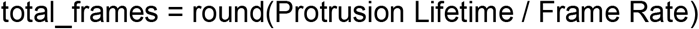

The protrusion window therefore corresponds to the interval:

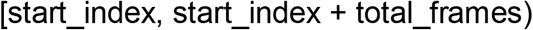

Directional parameters, curvature measurements, and NMII metrics were calculated strictly within this globally aligned protrusion window. Phase durations corresponding to expansion (E), stabilization (S), and retraction (R) were converted to frame counts and applied sequentially within the protrusion window to ensure consistent temporal alignment across cells. This coordinate-based slicing was used as the authoritative reference for all protrusion-resolved measurements to avoid inconsistencies arising from rounding differences between time-based and frame-based phase segmentation.

#### Principal component analysis and composite directionality index

To determine whether directional migration metrics reflected coordinated multivariate behavior rather than independent pairwise correlations, principal component analysis (PCA) was performed on all directional parameters measured for each protrusion. These parameters included net displacement (ND), total path length (TPL), straightness index (STA), persistence ratio (PR), angular variability (ANGVAR), turning index (TI), turning probability thresholds (TP15–TP85), and average speed.

Prior to PCA, all variables were standardized using z-score normalization:

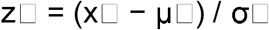

where x□ represents the observed value, μ□ the mean, and σ□ the standard deviation of variable i. Standardization ensured that all variables contributed equally to the analysis regardless of scale.

Principal components were computed using singular value decomposition. The first principal component (PC1) captured the dominant variance in directional behavior and was interpreted as the principal axis of directional persistence. For interpretability, the sign of PC1 was inverted when necessary so that higher PC1 values corresponded to stronger directional migration.

To generate a continuous metric of directional behavior, a Composite Directionality Index (CDI) was defined as the average of the standardized directional variables:

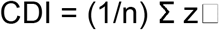

Protrusions were stratified into high- and low-directionality groups based on the median CDI value. Statistical comparisons between groups were performed using two-tailed tests as described in the statistical analysis section.

#### Temporal organization of protrusion states

To analyze how protrusions are temporally organized during migration, protrusions were first classified into high- and low-directionality states using the CDI-based classification described above. For each cell trajectory, protrusions were ordered according to their onset frame within the global coordinate series.

Transition probabilities between protrusion states were calculated by determining the frequency with which a protrusion of a given state was followed by another protrusion of the same or different state. Transition probabilities were computed as: P(A → B) = number of transitions from state A to state B / total number of transitions from state A

Run length was defined as the number of consecutive protrusions belonging to the same directional state. Mean run length was calculated separately for high- and low-directionality protrusion states within each cell trajectory.

These analyses allowed quantification of whether protrusion states occurred intermittently or formed extended sequences during migration.

#### Statistical analysis

All quantitative analyses were performed using MATLAB (MathWorks), GraphPad Prism (GraphPad Software), or custom scripts as described above. Data are presented as mean ± SEM unless otherwise indicated. For analyses comparing distributions of protrusion parameters or NMII metrics between groups, nonparametric two-tailed Mann–Whitney tests were used. For comparisons among multiple experimental conditions, one-way analysis of variance (ANOVA) followed by Tukey’s multiple-comparisons test was applied.

Correlation analyses between structural, directional, and NMII-related parameters were performed using Pearson correlation coefficients. Correlation matrices were computed using MATLAB, and correlation strengths are reported as Pearson r values. Heat maps representing correlation matrices display the magnitude and sign of Pearson coefficients.

For time-dependent measurements of NMII localization or other fluctuating cellular variables, threshold-based analyses were used to quantify the fraction of time individual cells maintained values above defined thresholds. Population differences across thresholds were evaluated using one-way ANOVA tests performed at each sampling point across independent experiments.

Principal component analysis (PCA) was performed on standardized directional parameters as described above to identify dominant axes of variation in protrusion behavior. A composite directionality index (CDI) was derived from the standardized directional variables and used to stratify protrusions or cells into high- and low-directionality groups based on median CDI values.

Unless otherwise stated, statistical tests were two-sided. Sample sizes (number of cells, protrusions, or independent experiments) are indicated in the corresponding figure legends. No statistical methods were used to predetermine sample size, but the numbers of observations analyzed are comparable to those commonly reported in the field. P values < 0.05 were considered statistically significant.

## References

1. de Oliveira, S., Rosowski, E. E. & Huttenlocher, A. Neutrophil migration in infection and wound repair: going forward in reverse. Nat. Rev. Immunol. 16, 378–391 (2016).

2. Liew, P. X. & Kubes, P. The Neutrophil’s Role During Health and Disease. Physiol. Rev. 99, 1223–1248 (2019).

3. Lämmermann, T. et al. Neutrophil swarms require LTB4 and integrins at sites of cell death in vivo. Nature 498, 371–375 (2013).

4. Mihlan, M., Safaiyan, S., Stecher, M., Paterson, N. & Lämmermann, T. Surprises from Intravital Imaging of the Innate Immune Response. Annu. Rev. Cell Dev. Biol. 38, 467–489 (2022).

5. Shaul, M. E. & Fridlender, Z. G. Tumour-associated neutrophils in patients with cancer. Nat. Rev. Clin. Oncol. 16, 601–620 (2019).

6. Wigerblad, G. & Kaplan, M. J. Neutrophil extracellular traps in systemic autoimmune and autoinflammatory diseases. Nat. Rev. Immunol. 23, 274–288 (2023).

7. SenGupta, S., Parent, C. A. & Bear, J. E. The principles of directed cell migration. Nat. Rev. Mol. Cell Biol. 22, 529 (2021).

8. Dorward, D. A. et al. The role of formylated peptides and formyl peptide receptor 1 in governing neutrophil function during acute inflammation. Am. J. Pathol. 185, 1172– 1184 (2015).

9. Weiner, O. D. Regulation of cell polarity during eukaryotic chemotaxis: the chemotactic compass. Curr. Opin. Cell Biol. 14, 196–202 (2002).

10. Saha, S., Town, J. P. & Weiner, O. D. Mechanosensitive mTORC2 independently coordinates leading and trailing edge polarity programs during neutrophil migration. Mol. Biol. Cell 34, ar35 (2023).

11. Shi, Y. et al. The mDial Formin Is Required for Neutrophil Polarization, Migration, and Activation of the LARG/RhoA/ROCK Signaling Axis during Chemotaxis. J. Immunol. 182, 3837–3845 (2009).

12. Xu, J. et al. Divergent signals and cytoskeletal assemblies regulate self-organizing polarity in neutrophils. Cell 114, 201–214 (2003).

13. Heit, B. & Kubes, P. Measuring Chemotaxis and Chemokinesis: The Under-Agarose Cell Migration Assay. Sci STKE 2003, pl5–pl5 (2003).

14. Lämmermann, T. et al. Rapid leukocyte migration by integrin-independent flowing and squeezing. Nature 453, 51–55 (2008).

15. Tsai, T. Y.-C. et al. Efficient Front-Rear Coupling in Neutrophil Chemotaxis by Dynamic Myosin II Localization. Dev. Cell 49, 189–205.e6 (2019).

16. Graziano, B. R. et al. Cell confinement reveals a branched-actin independent circuit for neutrophil polarity. PLOS Biol. 17, e3000457 (2019).

17. Li, J. L. et al. Intravital multiphoton imaging of immune responses in the mouse ear skin. Nat. Protoc. 7, 221–234 (2012).

18. Slaba, I. et al. Imaging the dynamic platelet-neutrophil response in sterile liver injury and repair in mice. Hepatol. Baltim. Md 62, 1593–1605 (2015).

19. Melis, N., Subramanian, B., Chen, D. & Weigert, R. Imaging Neutrophil Migration in the Mouse Skin to Investigate Subcellular Membrane Remodeling Under Physiological Conditions. J. Vis. Exp. JoVE https://doi.org/10.3791/63581 (2022) doi:10.3791/63581.

20. Barros-Becker, F., Lam, P.-Y., Fisher, R. & Huttenlocher, A. Live imaging reveals distinct modes of neutrophil and macrophage migration within interstitial tissues. J. Cell Sci. 130, 3801–3808 (2017).

21. Neupane, A. S. et al. Patrolling Alveolar Macrophages Conceal Bacteria from the Immune System to Maintain Homeostasis. Cell 183, 110–125.e11 (2020).

22. Choudhury, S. R. et al. Dipeptidase-1 Is an Adhesion Receptor for Neutrophil Recruitment in Lungs and Liver. Cell 178, 1205–1221.e17 (2019).

23. Maupin, P., Phillips, C. L., Adelstein, R. S. & Pollard, T. D. Differential localization of myosin-II isozymes in human cultured cells and blood cells. J. Cell Sci. 107 (Pt 11s), 3077–3090 (1994).

24. Golomb, E. et al. Identification and Characterization of Nonmuscle Myosin II-C, a New Member of the Myosin II Family. J. Biol. Chem. 279, 2800–2808 (2004).

25. Vicente-Manzanares, M., Ma, X., Adelstein, R. S. & Horwitz, A. R. Non-muscle myosin II takes centre stage in cell adhesion and migration. Nat. Rev. Mol. Cell Biol. 10, 778–790 (2009).

26. Zhang, Y. et al. Mouse models of MYH9-related disease: mutations in nonmuscle myosin II-A. Blood 119, 238–250 (2012).

27. Subramanian, B. C. et al. The LTB4-BLT1 axis regulates actomyosin and β2-integrin dynamics during neutrophil extravasation. J. Cell Biol. 219, e201910215 (2020).

28. Wong, K., Van Keymeulen, A. & Bourne, H. R. PDZRhoGEF and myosin II localize RhoA activity to the back of polarizing neutrophil-like cells. J. Cell Biol. 179, 1141– 1148 (2007).

29. Amornphimoltham, P., Thompson, J., Melis, N. & Weigert, R. Non-invasive intravital imaging of head and neck squamous cell carcinomas in live mice. Methods San Diego Calif 128, 3–11 (2017).

30. Calo, C. J., Patil, T., Palizzi, M., Wheeler, N. & Hind, L. E. Collagen concentration regulates neutrophil extravasation and migration in response to infection in an endothelium dependent manner. Front. Immunol. 15, 1405364 (2024).

31. Hind, L. E., Vincent, W. J. B. & Huttenlocher, A. Leading from the back: the role of the uropod in neutrophil polarization and migration. Dev. Cell 38, 161–169 (2016).

32. Saha, S., Town, J. P. & Weiner, O. D. Mechanosensitive mTORC2 independently coordinates leading and trailing edge polarity programs during neutrophil migration. Mol. Biol. Cell 34, ar35 (2023).

33. Aguilar-Cuenca, R. et al. Tyrosine Phosphorylation of the Myosin Regulatory Light Chain Controls Non-muscle Myosin II Assembly and Function in Migrating Cells. Curr. Biol. CB 30, 2446–2458.e6 (2020).

34. Li, Y. et al. PTENα promotes neutrophil chemotaxis through regulation of cell deformability. Blood 133, 2079–2089 (2019).

35. Mylvaganam, S., Freeman, S. A. & Grinstein, S. The cytoskeleton in phagocytosis and macropinocytosis. Curr. Biol. 31, R619–R632 (2021).

36. Insall, R. H. Understanding eukaryotic chemotaxis: a pseudopod-centred view. Nat. Rev. Mol. Cell Biol. 11, 453–458 (2010).

37. Niggli, V. Rho-kinase in human neutrophils: a role in signalling for myosin light chain phosphorylation and cell migration. FEBS Lett. 445, 69–72 (1999).

38. Vlahos, C. J., Matter, W. F., Hui, K. Y. & Brown, R. F. A specific inhibitor of phosphatidylinositol 3-kinase, 2-(4-morpholinyl)-8-phenyl-4H-1-benzopyran-4-one (LY294002). J. Biol. Chem. 269, 5241–5248 (1994).

39. Gambardella, L. & Vermeren, S. Molecular players in neutrophil chemotaxis--focus on PI3K and small GTPases. J. Leukoc. Biol. 94, 603–612 (2013).

40. Afonso, P. V. & Parent, C. A. PI3K and chemotaxis: a priming issue? Sci. Signal. 4, pe22 (2011).

41. Ebrahim, S. et al. Dynamic polyhedral actomyosin lattices remodel micron-scale curved membranes during exocytosis in live mice. Nat. Cell Biol. 21, 933–939 (2019).

42. Halder, D. et al. Nonmuscle myosin IIA and IIB differentially modulate migration and alter gene expression in primary mouse tumorigenic cells. Mol. Biol. Cell 30, 1463– 1476 (2019).

43. Shutova, M. S. et al. Self-sorting of nonmuscle myosins IIA and IIB polarizes the cytoskeleton and modulates cell motility. J. Cell Biol. 216, 2877–2889 (2017).

44. Driscoll, M. K. et al. Robust and automated detection of subcellular morphological motifs in 3D microscopy images. Nat. Methods 16, 1037–1044 (2019).

