## Supplementary Info for "Spatially Distinct Myosin II Architectures Regulate Protrusion Dynamics and Directional Persistence during Immune Cell Migration"

**Supplementary Figures**

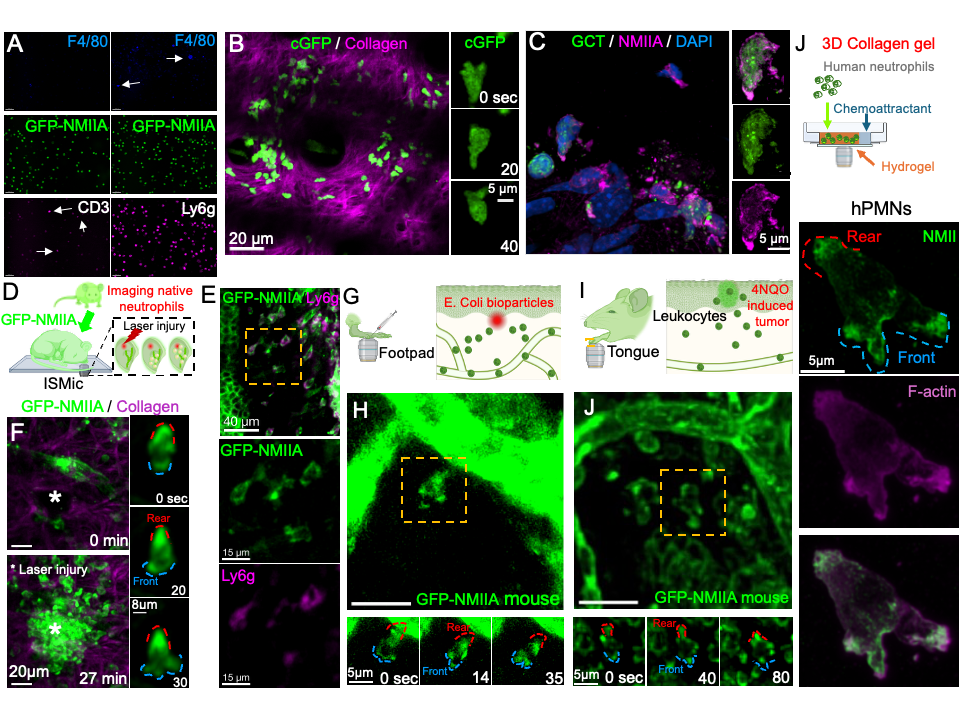

**Figure S1. NMIIA localizes at the front of migrating neutrophils in vivo and in three-dimensional environments.** (A) Immunofluorescence analysis of primary neutrophils isolated from GFP–NMIIA mice. Cells were fixed and labeled with immune cell markers to verify cell identity, including F4/80 and CD3 (left) or F4/80 and Ly6g (right). GFP–NMIIA fluorescence is shown in green. (B) Adoptive transfer control experiment using neutrophils expressing cytosolic GFP (cGFP). Maximum-intensity projections from two-photon intravital imaging of neutrophils migrating in the ear dermis following adoptive transfer into wild-type recipients. Collagen fibers are visualized by second harmonic generation (magenta). Time-lapse images of a representative cell show uniform cytosolic GFP distribution during migration. (C) Validation of NMIIA localization using wild-type neutrophils labeled ex vivo with green cell tracker (GCT) and immunostained for NMIIA after adoptive transfer. Maximum projections show cortical NMIIA localization in migrating neutrophils. (D) Schematic representation of the intravital imaging setup used to visualize native GFP–NMIIA neutrophils in the mouse ear during migration toward a laser-induced sterile injury. (E) Representative intravital two-photon images showing recruitment of native GFP–NMIIA neutrophils to the laser-injury site and their migration within the surrounding collagen network. Collagen fibers are visualized by second harmonic generation. (F) Fixed tissue from laser-injured ears immunostained for the neutrophil marker Ly6g (magenta), confirming the identity of GFP–NMIIA–positive cells. Insets show individual Ly6g⁺ neutrophils displaying NMIIA localization at both the front and rear cortex. (G) Schematic representation of the intravital imaging setup used to visualize neutrophil recruitment in the mouse footpad following injection of heat-inactivated *E. coli* bioparticles. (H) Two-photon imaging of GFP–NMIIA neutrophils recruited to the footpad, illustrating NMIIA localization during migration in vivo.(I) Schematic representation of the intravital imaging setup used to visualize neutrophil recruitment to carcinogen-induced tumors in the mouse tongue. (J) Two-photon imaging of GFP–NMIIA neutrophils migrating in the tumor microenvironment. Time-lapse insets in panels F, H, and J show representative migrating neutrophils with NMIIA localization along the cell cortex. Front and rear regions were defined based on the direction of migration and are indicated by blue and red dashed outlines, respectively.

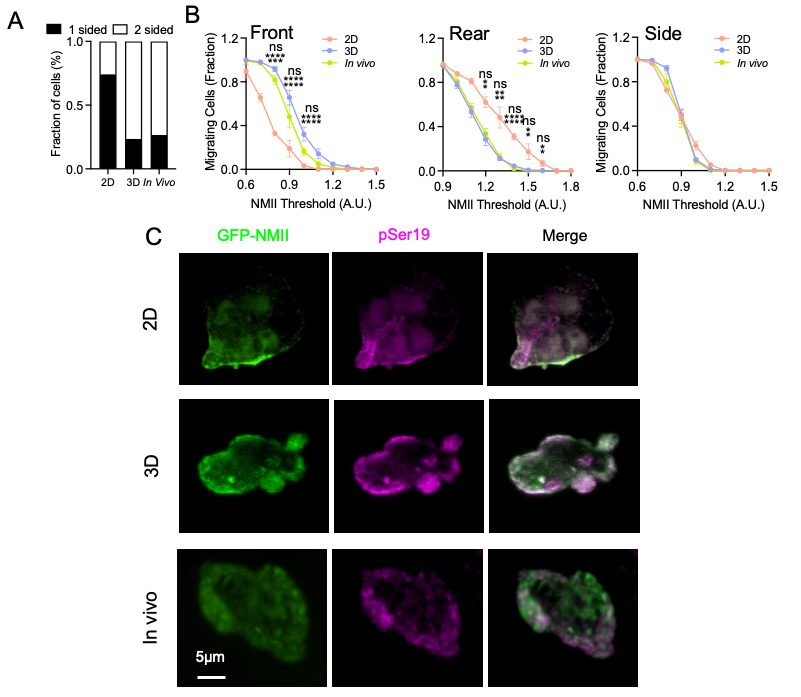

**Figure S2. Quantitative analysis of NMIIA distribution during neutrophil migration.**
(A) Distribution of NMII localization patterns in migrating neutrophils across environments. Cells were classified based on whether GFP–NMIIA was enriched at a single cortical region or at two opposing cortical regions of the cell. Quantification is shown as the fraction of cells displaying one-sided or two-sided NMII localization in 2D migration assays, 3D collagen matrices, and in vivo conditions (2D: 197 cells from 21 fields of view across 5 experiments; 3D: 286 cells from 10 fields of view across 3 experiments; in vivo: 384 cells from 7 fields of view across 4 animals).(B) Fraction of migrating cells maintaining NMII levels above increasing thresholds at different cortical regions. The proportion of cells maintaining NMII enrichment at the front, rear, or side of the cell for more than 67% of the migration time is plotted as a function of NMII intensity threshold for cells migrating in 2D, 3D collagen matrices, or in vivo. (C) Immunofluorescence detection of phosphorylated myosin regulatory light chain (pSer19) in GFP–NMIIA neutrophils migrating in 2D, 3D collagen matrices, or in vivo. Maximum intensity projections show GFP–NMIIA (green), phosphorylated MLC (pSer19, magenta), and merged images, indicating that NMIIA is present in an activated conformation across migration environments. Scale bar, 5 μm. Statistical comparisons were performed using one-way ANOVA with Tukey’s multiple comparisons test; significance levels are indicated in the figure.

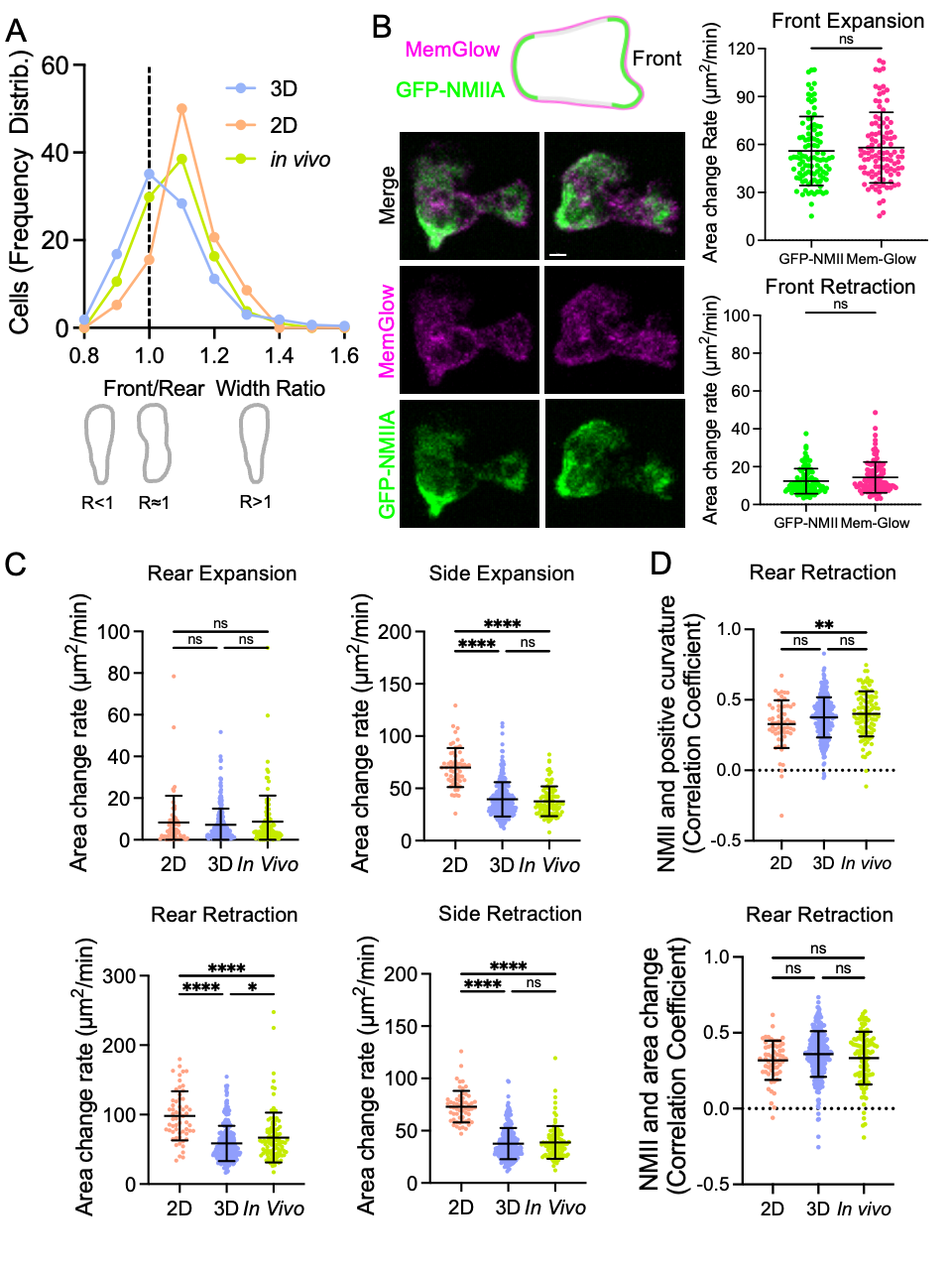

**Figure S3. Quantitative analysis of cell morphology, membrane dynamics, and NMII coupling during neutrophil migration.**(A) Distribution of the front-to-rear width ratio of migrating neutrophils in 2D migration assays, 3D collagen matrices, and in vivo conditions, illustrating differences in cell polarity across environments. Ratios >1 indicate a wider front relative to the rear, whereas ratios <1 indicate a wider rear.
(B) Validation of membrane boundary measurements using an independent membrane marker. Neutrophils expressing GFP–NMIIA were labeled with the membrane dye MemGlow to visualize the plasma membrane independently of NMII signal. Representative images show GFP–NMIIA (green), MemGlow (magenta), and merged channels. Quantification of membrane expansion and retraction rates measured using MemGlow confirms that membrane dynamics derived from GFP–NMIIA–based segmentation accurately reflect bona fide membrane behavior. (C) Quantification of expansion and retraction rates at the rear and lateral regions of migrating neutrophils across migration environments (2D, 3D collagen gels, and in vivo). (D) Correlation analysis between NMII localization and membrane remodeling dynamics at the rear cortex, including correlations between NMII levels and positive membrane curvature or local area changes associated with retraction. Data are shown as mean ± SEM across independent experiments (N = 3, 11, and 5 for 2D, 3D, and in vivo conditions, respectively; total number of single-cell trajectories n = 58, 268, and 104). Statistical comparisons were performed using one-way ANOVA with Tukey’s multiple comparisons test; significance levels are indicated in the figure.

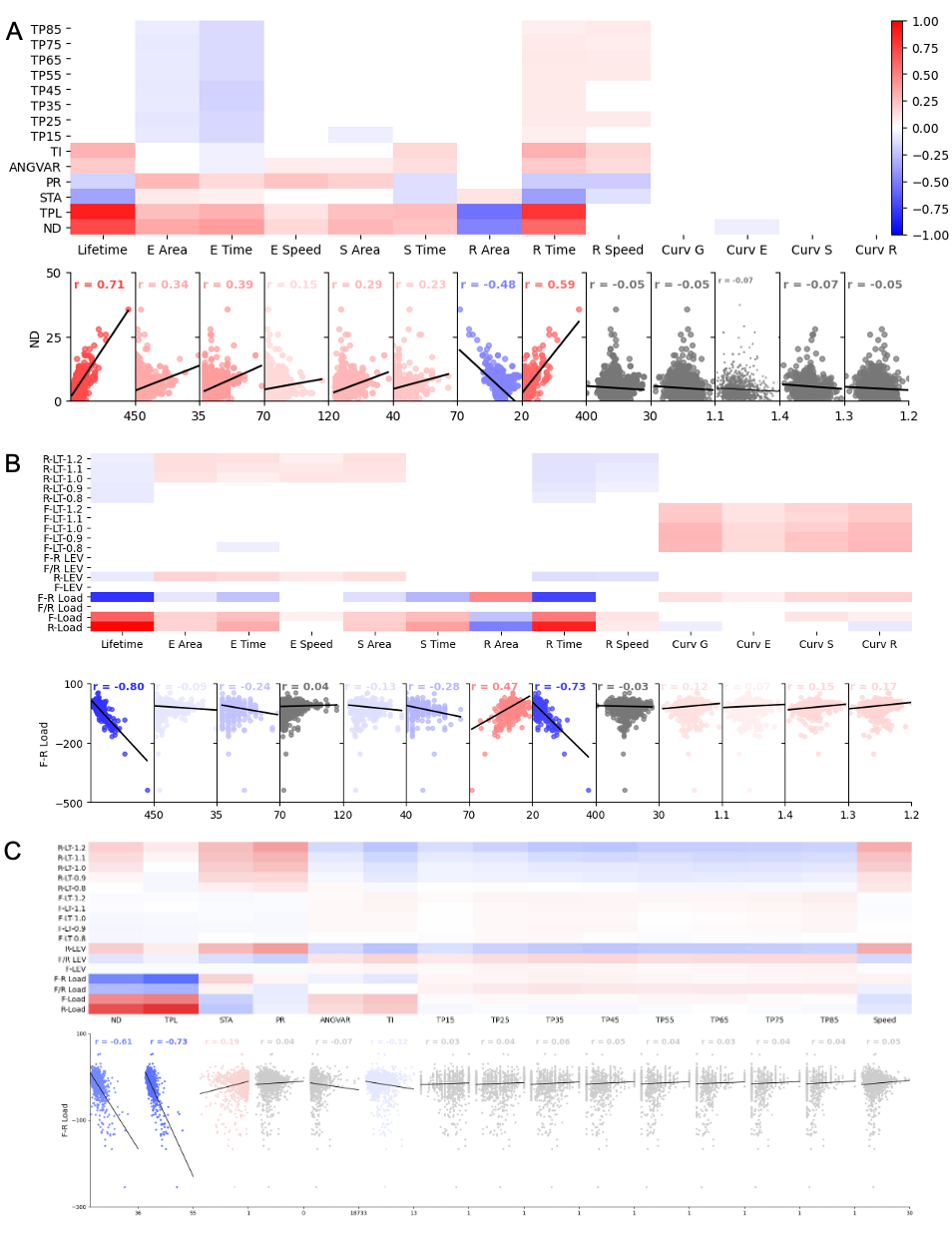

**Figure S4. Extended correlation analyses linking protrusion structure, NMII engagement, and directional migration.** (A) Relationships between protrusion structural parameters and directional migration. Upper panel: heatmap showing Pearson correlation coefficients between protrusion structural features (lifetime, phase durations, phase areas, phase speeds, and curvature metrics) and directional parameters including net displacement (ND), total path length (TPL), straightness (STA), persistence ratio (PR), angular variability (ANGVAR), turning index (TI), and turning probability thresholds (TP15–TP85). Lower panels show representative scatter plots illustrating correlations between ND and selected protrusion structural parameters. (B) Relationships between NMII engagement and protrusion structural parameters. Upper panels show heatmaps of Pearson correlations between NMII metrics—including rear NMII load, front NMII load, front–rear NMII load difference (FR load), front/rear ratio, and threshold-based NMII lifetime parameters—and protrusion structural features. Lower panels show representative scatter plots illustrating correlations between FR load and selected protrusion structural parameters. (C) Relationships between NMII engagement and directional migration parameters. Heatmap and scatter plots show correlations between FR load and directional migration metrics (ND, TPL, STA, PR, ANGVAR, TI, TP15–TP85, and migration speed). Rear-biased NMII engagement is associated with increased directional persistence and inversely correlated with turning-related parameters.

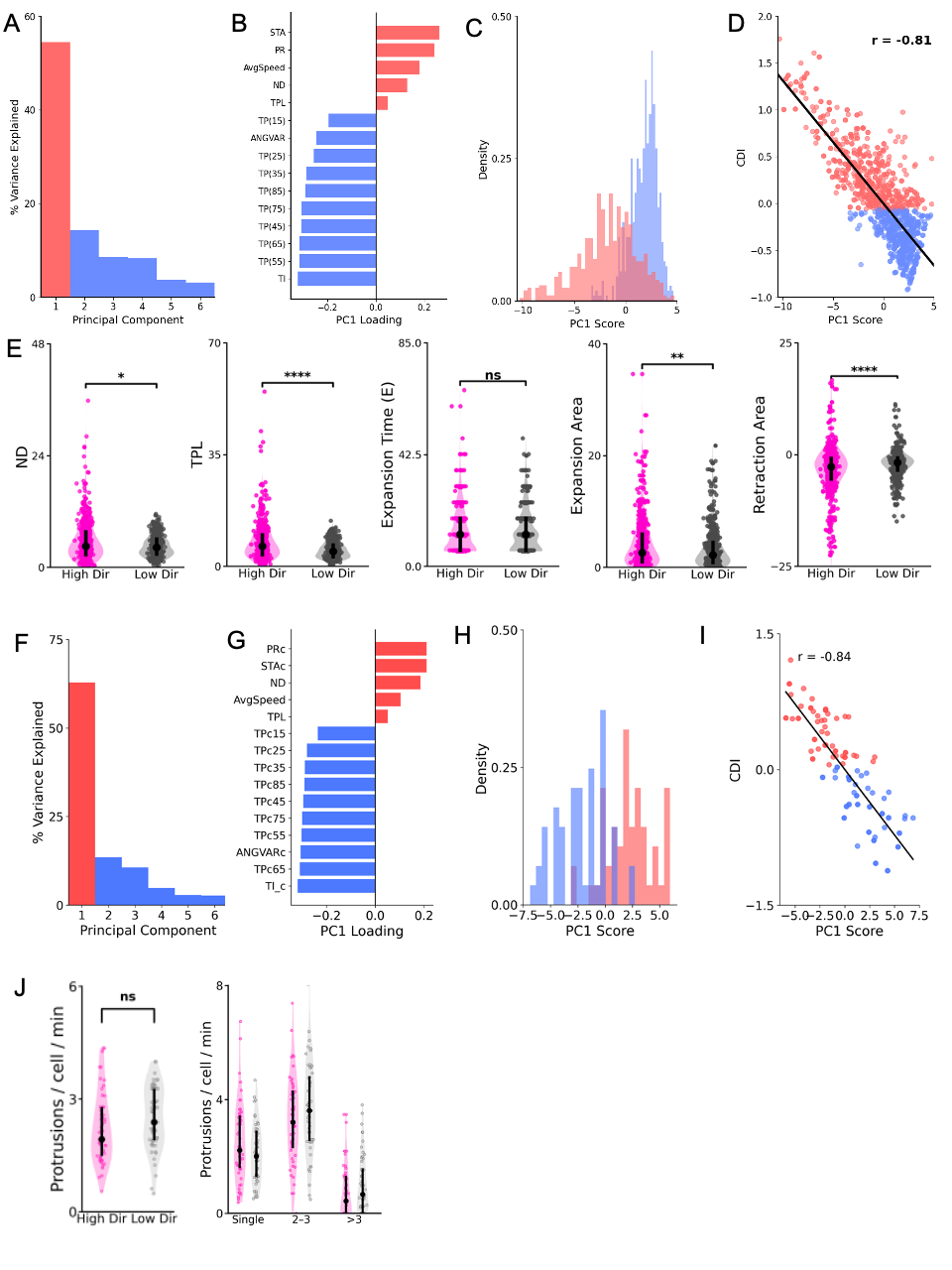

**Figure S5. Principal component and composite directionality analyses.**
(A) Variance explained by principal components derived from protrusion directional parameters. The first principal component (PC1) captures the dominant variance associated with directional persistence.
(B) Loadings of directional parameters on PC1, indicating the relative contribution of migration metrics including straightness (STA), persistence ratio (PR), average speed, net displacement (ND), total path length (TPL), angular variability (ANGVAR), turning index (TI), and turning probability thresholds (TP15–TP85). (C) Distribution of PC1 scores across protrusions(D) Relationship between PC1 scores and the composite directionality index (CDI), showing strong correspondence between PCA-derived and composite directionality metrics. (E) Comparison of structural protrusion parameters between high- and low-directionality protrusions, including net displacement (ND), total path length (TPL), expansion duration, expansion area, and retraction area. Statistical comparisons were performed using Mann–Whitney tests. (F) Variance explained by principal components derived from cell-level directional parameters. (G) Loadings of directional parameters contributing to PC1 at the cell level. (H) Distribution of PC1 scores across cells.( I) Relationship between PC1 scores and CDI at the cell level. (J) Analysis of protrusion production across cells classified as high- or low-directionality, including protrusion frequency per cell and distribution of cells producing single, two to three, or more than three protrusions.

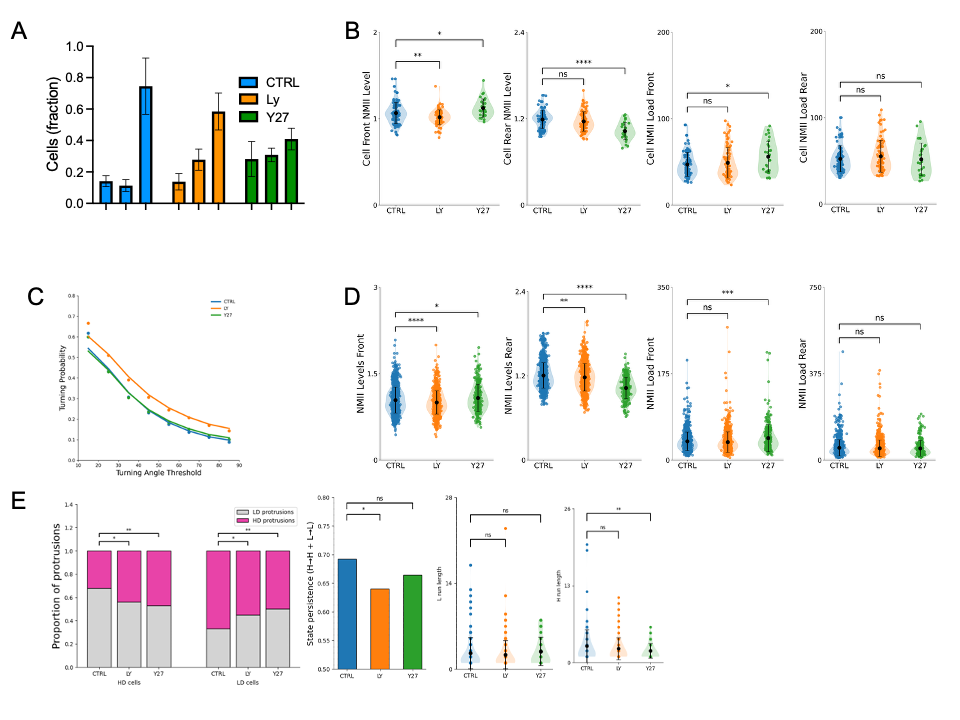

**Figure S6. Extended analyses of pharmacological perturbations affecting NMII organization and protrusion dynamics.** (A) Optimization of pharmacological perturbation conditions. The concentrations of ROCK inhibitor Y27632 and PI3K inhibitor LY294002 used in this study were selected to partially perturb NMII organization while preserving cell motility. Quantification shows the fraction of motile cells under control and inhibitor-treated conditions. (B) Extended analysis of NMII persistence across intensity thresholds. The fraction of migrating cells maintaining NMII levels above increasing thresholds at the cell front or rear is plotted for control, LY294002, and Y27632 conditions, revealing reduced front NMII persistence after PI3K inhibition and reduced rear NMII persistence following ROCK inhibition.(C) Distributions of turning probability across angular thresholds. PI3K inhibition increases turning behavior across thresholds, whereas ROCK inhibition primarily affects net displacement with comparatively modest effects on turning probability. (D) Structural protrusion parameters across pharmacological conditions, including protrusion lifetime, phase durations, protrusion areas, expansion speed, and curvature metrics. Protrusion lifetimes remain largely unchanged across perturbations, whereas PI3K inhibition shortens the stabilization phase and ROCK inhibition reduces protrusion expansion speed. (E) Phase-resolved analysis of NMII engagement and directional metrics. Measurements of NMII lifetime, angular variability, and turning parameters during protrusion expansion, stabilization, and retraction phases reveal polarity-specific effects of the inhibitors, with PI3K inhibition destabilizing front-associated NMII during expansion and ROCK inhibition reducing rear NMII persistence during retraction. Statistical comparisons were performed using the tests indicated in the figure.

**Legends to Supplementary Movies**

**Movie S1. Migration of neutrophils expressing GFP-NMII in the mouse ear (Adoptive transfer) - Low magnification.**

Adoptive transfer 2-photon imaging of primary neutrophils isolated from a GFP-NMIIA mouse bone marrow injected in the ear of a recipient wt mouse. * Sterile injury; Green: GFP-NMIIA; Purple: Collagen-I SHG.

**Movie S2. Migration of neutrophils expressing GFP-NMII in the mouse ear (Adoptive transfer) - High magnification.**

Adoptive transfer 2-photon imaging of primary neutrophils isolated from a GFP-NMIIA mouse bone marrow injected in the ear of a recipient wt mouse. Green: GFP-NMIIA.

**Movie S3. Migration of endogenous neutrophils in the ear of a GFP-NMIl expressing mouse (Direct imaging) - Low magnification.**

2 photon direct imaging of native neutrophils in the ear of a GFP-NMIIA mouse after a sterile laser injury. * Sterile injury; Green: GFP-NMIIA; Purple: Collagen-I SHG.

**Movie S4. Migration of endogenous neutrophils in the ear of a GFP-NMIl expressing mouse (Direct imaging) - High magnification.**

2 photon direct imaging of native neutrophils in the ear of a GFP-NMIIA mouse after a sterile laser injury. Green: GFP-NMIIA

**Movie S5. Migration of endogenous neutrophils in the footpad of a GFP-NMIl expressing mouse in response to E. coli bioparticles (Direct imaging).**

2 photon direct imaging of native neutrophils in the hind footpad of a GFP-NMIIA mouse after injection of heat inactivated E. coli bioparticles. Green: GFP-NMIIA; Purple: Collagen-I SHG.

**Movie S6. Migration of endogenous neutrophils in the tongue of a GFP-NMIl expressing mouse in the vicinity of a tumor (Direct imaging).**

2 photon direct imaging of native neutrophils in the tongue of a GFP-NMIIA mouse after carcinogen (4NQO) induced tumor. Green: GFP-NMIIA; Purple: Collagen-I SHG.

**Movie S7. Migration of neutrophils expressing GFP-NMII in 3D (collagen-I hydrogel).**

2 photon imaging of primary neutrophils isolated from a GFP-NMIIA mouse bone marrow embedded in a 3D collagen hydrogel. Cells are migrating in response to a chemoattractant (WKYMVm, 0.3 µM final) stimulation. Green: GFP-NMIIA.

**Movie S8. Migration of neutrophils expressing GFP-NMII in 2D (Under-Agarose).**

2 photon imaging of primary neutrophils isolated from a GFP-NMIIA mouse bone marrow in a 2D under-agarose migration assay. Cells are seeded in a well and migration is triggered by a chemoattractant gradient (WKYMVm, 0.3 µM final). Green: GFP-NMIIA.

**Movie S9. Migration of neutrophils expressing GFP-NMII in 3D treated with a ROCK inhibitor (Y27632).**

2 photon imaging of primary neutrophils isolated from a GFP-NMIIA mouse bone marrow embedded in a 3D collagen hydrogel. Cells are pre-treated with 30µM of Y27632 before migrating in response to a chemoattractant (WKYMVm, 0.3 µM final) stimulation. Green: GFP-NMIIA.

**Movie S10. Migration of neutrophils expressing GFP-NMII in 3D treated with a PI3K inhibitor (Ly294002).**

2 photon imaging of primary neutrophils isolated from a GFP-NMIIA mouse bone marrow embedded in a 3D collagen hydrogel. Cells are pre-treated with 10µM of Ly294002 before migrating in response to a chemoattractant (WKYMVm, 0.3 µM final) stimulation. Green: GFP-NMIIA.
